# Readthrough errors purge deleterious cryptic sequences, facilitating the birth of coding sequences

**DOI:** 10.1101/737452

**Authors:** Luke Kosinski, Joanna Masel

## Abstract

*De novo* protein-coding innovations sometimes emerge from ancestrally non-coding DNA, despite the expectation that translating random sequences is overwhelmingly likely to be deleterious. The “pre-adapting selection” hypothesis claims that emergence is facilitated by prior, low-level translation of non-coding sequences via molecular errors. It predicts that selection on polypeptides translated only in error is strong enough to matter, and is strongest when erroneous expression is high. To test this hypothesis, we examined non-coding sequences located downstream of stop codons (i.e. those potentially translated by readthrough errors) in *Saccharomyces cerevisiae* genes. We identified a class of “fragile” proteins under strong selection to reduce readthrough, which are unlikely substrates for co-option. Among the remainder, sequences showing evidence of readthrough translation, as assessed by ribosome profiling, encoded C-terminal extensions with higher intrinsic structural disorder, supporting the pre-adapting selection hypothesis. The cryptic sequences beyond the stop codon, rather than spillover effects from the regular C-termini, are primarily responsible for the higher disorder. Results are robust to controlling for the fact that stronger selection also reduces the length of C-terminal extensions. These findings indicate that selection acts on 3′ UTRs in *S. cerevisiae* to purge potentially deleterious variants of cryptic polypeptides, acting more strongly in genes that experience more readthrough errors.

## Introduction

One of the more surprising recent developments in molecular evolution is that adaptive protein-coding innovations sometimes arise out of DNA sequences that were previously non-coding. Sometimes only part of a protein arises in this way, via new or expanded coding exons (Sorek 2007), or via annexation of 3′ untranslated regions (UTRs) (Giacomelli, et al. 2007; Vakhrusheva, et al. 2011; Andreatta, et al. 2015) or 5′ UTRs (Wilder, et al. 2009) into an ORF. More dramatically, completely new protein-coding genes can also arise *de novo* (reviewed in McLysaght and Guerzoni 2015; Van Oss and Carvunis 2019). This contradicts the classic stance that it is implausible under modern conditions of life for a useful protein to appear *de novo* from a random sequence of nucleotides with no history of selection (Zuckerkandl 1975; Jacob 1977).

As an example of how important a history of selection is for protein function, consider amino acid composition. Hydrophilic residues promote intrinsic structural disorder (ISD), i.e. a propensity to avoid stable conformational structures and instead exist as a dynamic ensemble of many conformational sub-states separated by low energy barriers (Guharoy, et al. 2015). Disordered regions are a key component of many proteins and perform important functions such as scaffolding and DNA binding (Uversky 2013; Habchi, et al. 2014; Guharoy, et al. 2015). Disordered proteins are also less prone to toxic aggregation (Linding, et al. 2004; Angyan, et al. 2012). Proteins are more disordered than would be expected from the translation of intergenic (Wilson, et al. 2017) or frameshifted (Willis and Masel 2018) sequences, as a consequence of a history of selection on their protein products.

The *de novo* evolution of a protein-coding sequence occurs when a “co-option” mutation converts a non-coding sequence into a coding sequence, e.g. a point mutation destroys a stop codon, or a mutation makes an intergenic ORF more prone to translation. Prior to such a co-option mutation taking place, we call the hypothetical gene product a “potential polypeptide.” Naïvely, we do not expect potential polypeptides to have been previously exposed to the selection needed to purge dangerous (e.g. excessively hydrophobic) sequence variants. However, this is not a foregone conclusion, because potential polypeptides are translated at low levels, as a consequence of widespread spurious transcription (Clark, et al. 2011; Tisseur, et al. 2011; Palazzo and Lee 2015; Neme and Tautz 2016; Blevins, et al. 2019) and translation (Wilson and Masel 2011; Ruiz-Orera, et al. 2014; Ji, et al. 2015; Ruiz-Orera, et al. 2018; Blevins, et al. 2019; Durand, et al. 2019).

The products of this spurious translation of non-coding sequences provide a preview of the effects of possible future co-option mutations (fig. 1). To show how, consider the case of reading beyond a stop codon. When a ribosome skips a stop codon, the same sequence beyond the stop codon is translated as would become the norm were a mutation to destroy or frameshift that stop codon. When this 3′ UTR sequence contains a backup stop codon, a stop codon readthrough error and a stop codon mutation will append the same C-terminal extension to a protein (fig. 1A and 1B). If no backup stop codon exists, both will trigger non-stop mediated mRNA decay (Vasudevan, et al. 2002) (fig. 1C and 1D). If a backup stop codon exists, but the extension sequence is unfavorable, the protein may be targeted for degradation (Arribere, et al. 2016). In all three cases, we refer to the sequence beyond the stop codon as a “cryptic sequence.” The cryptic sequence encodes both the consequences of expressing the potential polypeptide via readthrough, and the consequences of a future stop codon loss mutation. Normally, we consider selection as always occurring after mutation. But because a stop codon readthrough error previews a future stop codon mutation, selection on the consequences of a mutation can act before mutation occurs.

**Fig. 1.**
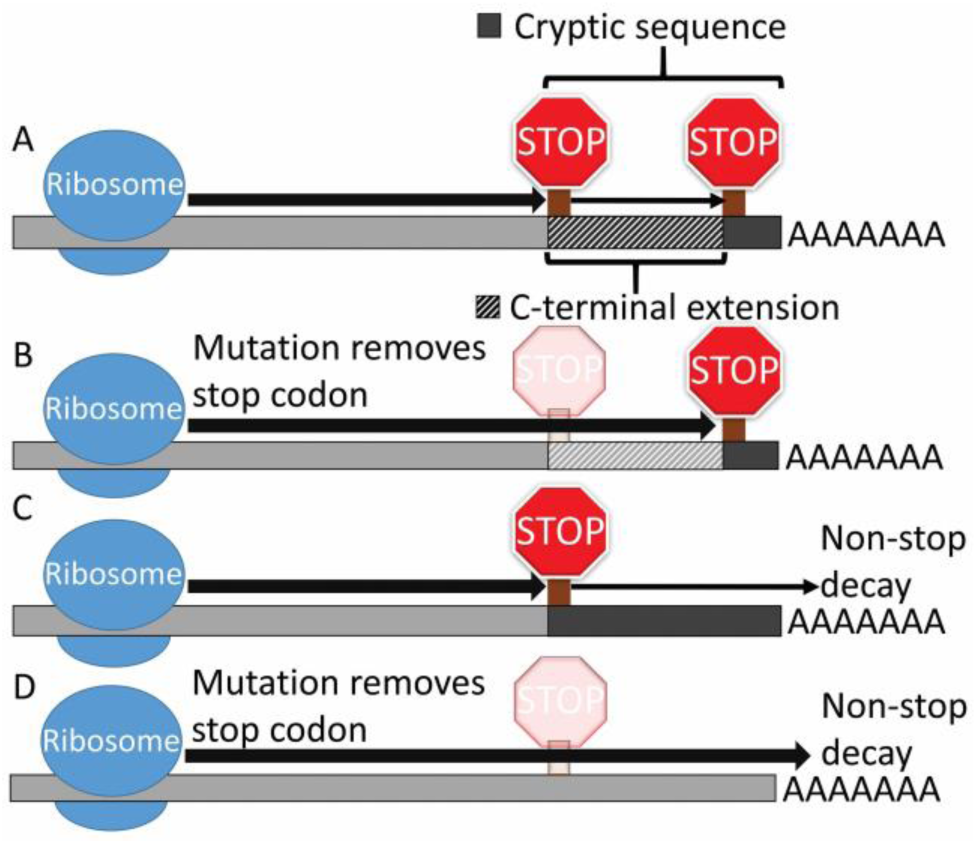
Stop codon readthrough previews the effects of stop codon mutations. Large and small black arrows represent constitutive and readthrough-dependent translation, respectively. When a backup stop codon exists (A and B), both translate the cryptic sequence between the stop codon and the backup stop codon, appending the product to the protein as a C-terminal extension. If there is no backup stop codon before the end of the 3’UTR (C and D), translation proceeds into the poly A tail, triggering non-stop decay instead of producing a C-terminal extended protein product.

This similarity between the consequences of a gene expression “error” in the present and a mutation in the future is not restricted to stop codon loss. For example, splicing errors/splice site mutations can include an intron in an existing ORF, and near-cognate start codons can easily mutate to become constitutive start codons. More broadly, mutations might also convert a promiscuous enzymatic or binding interaction into a constitutive one (Khersonsky and Tawfik 2010; Pal and Papp 2017). While we focus here on stop codon readthrough, the principles we illustrate are broadly applicable.

We expect the distribution of fitness effects (DFE) of errors to be a dampened version of the corresponding DFE of new co-option mutations, because translation errors lead to the same expression or degradation events, at lower levels. In general, the DFE of new mutations is bimodal, with strongly deleterious mutations forming one mode and relatively benign mutations (mostly weakly deleterious and neutral, but also including rare beneficial mutations) forming the other (Keightley and Eyre-Walker 2007; Lind, et al. 2017). Intuitively, these two modes correspond, in the case of changes to an existing protein, to mutations that break their associated protein and are strongly deleterious, versus those that make a minor tweak and are relatively “benign.” We consider potential polypeptides to be “benign” or “deleterious” according to the DFE mode of their corresponding co-option mutation. In order for a co-option mutation to contribute to evolutionary innovation, it must not cause too much collateral harm, i.e. it must co-opt a benign potential polypeptide (Masel 2006; Rajon and Masel 2011).

Some proteins may be resilient to mutations adding amino acids to their C-terminal, while other proteins may be “fragile,” corresponding to significant vs. negligible frequencies for the benign mode of the DFE of possible C-terminal extensions. For example, proteins with C-terminal ends that are ordered and buried within the folds of the protein structure, making an extension likely to disrupt folding, may be particularly fragile to stop codon mutations.

If the DFE of translation errors is a dampened version of the DFE of new co-option mutations, then erroneous translation of benign potential polypeptides will be effectively neutral, and translation of strongly deleterious potential polypeptides will be only weakly deleterious, with the magnitude of deleterious-ness dependent on the error rate (Masel 2006; Rajon and Masel 2011). A mutation to a cryptic sequence that converts its erroneous expression from benign to deleterious will be effectively purged if and only if the deleterious effect of the corresponding potential polypeptide is strong enough and its translation by error is frequent enough (Xiong, et al. 2017). In other words, if errors occur frequently enough, potential polypeptides of strong effect will be exposed to enough selection to remove cryptic sequences with dangerous effects, leaving behind benign cryptic sequences that form better raw material for *de novo* innovations (Rajon and Masel 2011; Wilson and Masel 2011; Xiong, et al. 2017).

We use the term “pre-adapting” selection to describe selection that purges deleterious potential polypeptides before a co-option mutation occurs. The goal of this paper is to investigate whether pre-adapting selection occurs in nature. The pre-adapting selection hypothesis has the potential to explain how *de novo* protein-coding sequences avoid toxicity and hence can sometimes be adaptive, by showing that they have a history of exposure to and purging by selection (Wilson and Masel 2011).

Pre-adapting selection theory predicts that selective pressures on potential polypeptides are sometimes strong enough to purge deleterious variants, especially in populations of large effective population size. As a more testable corollary within a single genome, potential polypeptides expressed at higher levels are predicted to be more likely to be benign (Xiong, et al. 2017). To test this, we need both to quantify relative levels of erroneous translation of different potential polypeptides, and to identify correlates of polypeptide benign-ness, in order to show an association between the two.

To measure erroneous translation, we focus on stop codon readthrough errors that preview *de novo* C-terminal extensions to existing genes. While the translation of intergenic ORFs is perhaps of more interest because it previews complete *de novo* genes, *de novo* C-terminal extensions are more tractable while still being relevant to *de novo* protein-coding evolution. Erroneous translation products, whether from an intergenic ORF or from part of a 3′ UTR, are difficult to detect by proteomic methods because they tend to be short and sparse, and may be targeted by degradation machinery (Arribere, et al. 2016). We therefore quantify translation through ribosomal association with transcripts. This is not straightforward for intergenic ORFs (Ruiz-Orera, et al. 2018; Durand, et al. 2019), e.g. transcriptional analysis must confirm that translation is contiguous, and unannotated functional proteins must be excluded. In contrast, cryptic sequences past the stop codon are not part of the functional protein (see paragraph below) and are already known to be contiguously transcribed. It is therefore straightforward to count ribosome profiling hits (“ribohits”) beyond the stop codon as a direct measure of readthrough. This is better than using protein abundance as an indirect measure of opportunities for readthrough, because highly abundant proteins tend to have lower readthrough rates per translation event (Bonetti, et al. 1995; Li and Zhang 2019). Our ability to directly quantify the amount of erroneous translation using ribohits makes stop codon readthrough a perfect test case for assessing whether pre-adapting selection can ever occur.

One possible concern with using ribohits is that not all stop codon readthrough is necessarily in error. For example, *Drosophila melanogaster* has been shown to have significant functional (“programmed”) readthrough (Jungreis, et al. 2011). However, pervasive programmed readthrough seems to be phylogenetically restricted to Pancrustaceae (Jungreis, et al. 2016), and does not occur in our focal species of *S. cerevisiae* (Jungreis, et al. 2016; Li and Zhang 2019). We can therefore assume that ribohits to 3’ UTR sequences indicate translational errors, rather than alternative already-functional protein products. Some ribohits might also indicate non-translating ribosomes, especially in mutant yeast (Guydosh and Green 2014; Young, et al. 2015; Yordanova, et al. 2018); appropriate ribosome profiling methodology (Gerashchenko and Gladyshev 2014; Miettinen and Björklund 2015) in wild-type yeast can reduce the scale of this problem.

We expect benign polypeptides to share some traits, such as a tendency towards high ISD, with functional proteins that have been shaped by selection. Once such traits are identified, they can be used to differentiate potential polypeptides that are likely to be benign (and hence co-optable, as has been documented to occur (Giacomelli, et al. 2007; Vakhrusheva, et al. 2011; Andreatta, et al. 2015)) from those that are likely to be deleterious. We call these traits “preadaptations” because they are found prior to co-option and facilitate future adaptation. Longer extensions are more likely to do harm; this is reflected in their tendency to be degraded (Arribere, et al. 2016). Selection on cryptic sequences should therefore shorten potential polypeptides. However, while documenting selection for short selection length can demonstrate that selection is powerful enough to act, shortness is not the most interesting benign trait with respect to potential to contribute to substantial (i.e. longer) protein-coding innovations. Our primary focus is therefore on high ISD as a preadaptation.

The term “preadaptation” has a complicated history, so some discussion is warranted to avoid conflating our use with the various past uses of the term. Cuénot (1914) discussed “preadapted” characters that are either neutral or adaptive in one environment, then fortuitously become useful in a later environment (are co-opted) (Casinos 2017). Gould and Vrba (1982) took issue with this term, claiming it was implicitly teleological despite Cuénot’s explicit rejection of orthogenesis, and attempted to replace “preadaptation” with the mostly-synonymous term “exaptation” (Casinos 2017). Gould and Vrba (1982) distinguished between two types of exaptation, depending on whether co-option is of a character previously shaped by natural selection for a different function (an adaptation), versus a character not shaped by natural selection for any particular function (a nonaptation). Potential polypeptides are nonaptations.

Gould and Vrba (1982) did not consider systematic variation among nonaptations in their probabilities that future co-option might be beneficial. In contrast, Eshel and Matessi (1998) argued that there may be systematic variation in suitability for future adaptive co-option. Such variation justifies the use of the prefix “pre-”, even in the absence of a teleological claim. Specifically, Eshel and Matessi (1998) used the term “preadaptation” to refer to the presence of cryptic genetic variants (i.e., co-optable nonaptations) whose effects are enriched for trait values with an elevated probability of being adaptive after an environmental shift. Populations become preadapted in this sense because future environmental changes tend to resemble existing marginal environments (Eshel and Matessi 1998).

Masel (2006) similarly justified the use of the term “preadaptation” to describe a somewhat different process that also leads to systematic variation in suitability for co-option. When a stock of cryptic variants is expressed at low levels, it can be purged of variants in the deleterious mode of the DFE (which have no chance of becoming adaptive exaptations). As discussed above, this enriches, via a process of elimination, for variants in the benign mode, which have more adaptive potential. This process, which we here call “pre-adapting selection,” occurs when cryptic variants are not completely phenotypically silent, but are instead expressed at low levels (Masel 2006; Rajon and Masel 2011). Low expression of cryptic variants can also help cross fitness valleys (Whitehead, et al. 2008).

Here we follow Wilson et al. (2017) in using the term “preadaptation” to refer not to the process described by Masel (2006), but to a trait that systematically makes a protein less likely to be harmful. Our usage here thus distinguishes clearly for the first time between the process of “pre-adapting selection” and the existence of traits that are “preadaptations.” This distinction is necessary, because pre-adapting selection may be occurring even if the nature of preadaptations is unknown, while preadaptations might be identifiable even if pre-adapting selection is not in force. Preadaptation can simply refer to backward-time conditional probability; given that *de novo* gene birth occurred, new genes are more likely than the average non-coding sequence to have properties that facilitate gene birth.

We hypothesize that cryptic sequences are subject to pre-adapting selection. Cryptic sequences with a history of higher levels of pre-adapting selection (e.g. due to higher expression) should therefore display more preadaptation, i.e. they should be more likely to be benign. High ISD/hydrophilicity is a preadaptation for sequences beyond stop codons (Arribere, et al. 2016), just as it is for the *de novo* birth of complete genes (Wilson, et al. 2017; Willis and Masel 2018). We test whether sequences beyond yeast stop codons that show more evidence of translation, as indicated by ribosomal profiling, encode higher ISD.

## Results

### Likely targets of non-stop decay have no readthrough ribohits

Fragile proteins are those which are prone to breaking when a C-terminal extension is added. Their C-terminal extensions are therefore unlikely to contribute to *de novo* evolution. Our ability to detect evolvability-relevant pre-adapting selection will be enhanced if we are able to identify and exclude the most fragile proteins.

Proteins that are fragile to readthrough are expected to be under strong selection for low readthrough rates. In the rare cases where a fragile protein is read through, degrading the readthrough product can mitigate its harms. If a 3′ UTR has no backup stop codon, readthrough should trigger non-stop mRNA decay, degrading the readthrough product and the potentially faulty mRNA that produced it (Vasudevan, et al. 2002). We therefore expect selection on fragile proteins to remove backup stop codons from their 3’UTRs, as well as to lower their readthrough rates.

We tested whether lacking a backup stop codon is an indicator of low readthrough. There are three distinct frames that might have or lack a backup stop codon. Readthrough can either be in-frame (where the stop codon is read by a near-cognate tRNA) or result from a frameshift to one of two different frames. Genes lacking a backup stop codon in at least one frame do not contain any ribohits in their 3′ UTRs (fig. 2, P < 10^−300^, Pearson’s Chi-squared test on contingency table of presence/absence).

**Fig. 2.**
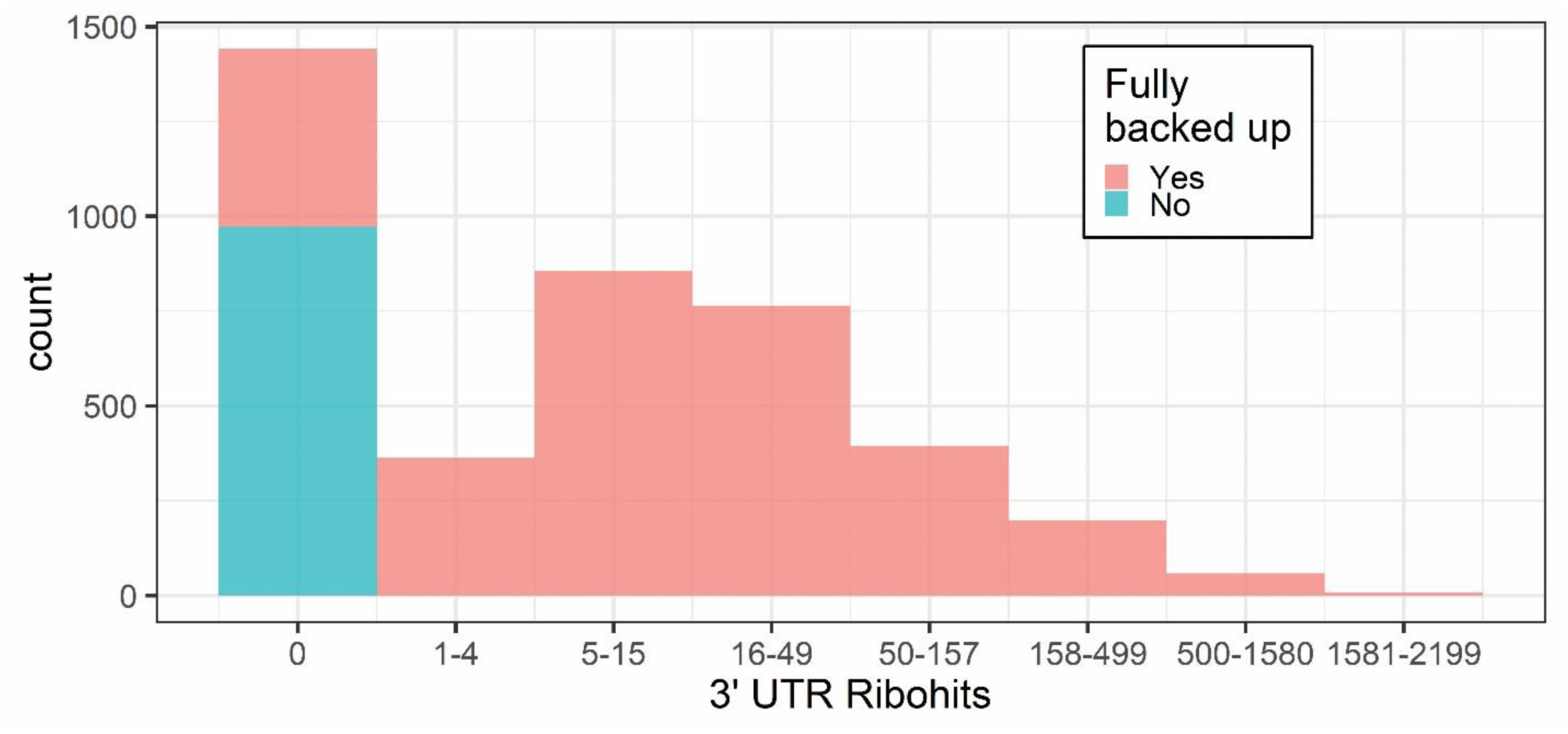
Histogram of ribohits coverage for 3’ UTRs in the Jarosz et al. (manuscript in preparation) dataset. 3’ UTR ribohits are divided into bins of width 0.5 on a log_10_ scale.

The dramatically lower levels of ribohits in genes lacking a backup stop codon cannot be explained by fewer opportunities to bind due to lower abundance, shorter 3’ UTRs, or shorter extensions in frames that still have a backup stop codon. Using a uniform [0, 1] prior, we obtain 95% credibility intervals for the probability of seeing at least one ribohit of 0.88-0.90 when all 3 frames have a backup stop, and 0-0.003 otherwise (see code on GitHub). We are thus fairly confident of at least a 290-fold difference between the two groups – and this is conservative, in neglecting the fact that most extensions with at least one ribohit have many. In contrast, in genes in which a least one frame lacks a backup stop codon, protein abundance is 1.4-fold lower (P = 3 × 10^−6^, Student’s t-test), 3’ UTRs are about 2-fold shorter (fig. 3A), and distances to the next backup stop codon (in frames where one exists) are 1.2 to 1.3-fold shorter, depending on frame (fig. 3B), with considerable overlap between the distributions (supplementary fig. 1). These modest differences in opportunities to bind fall far short of explaining a 300-fold difference.

**Fig. 3.**
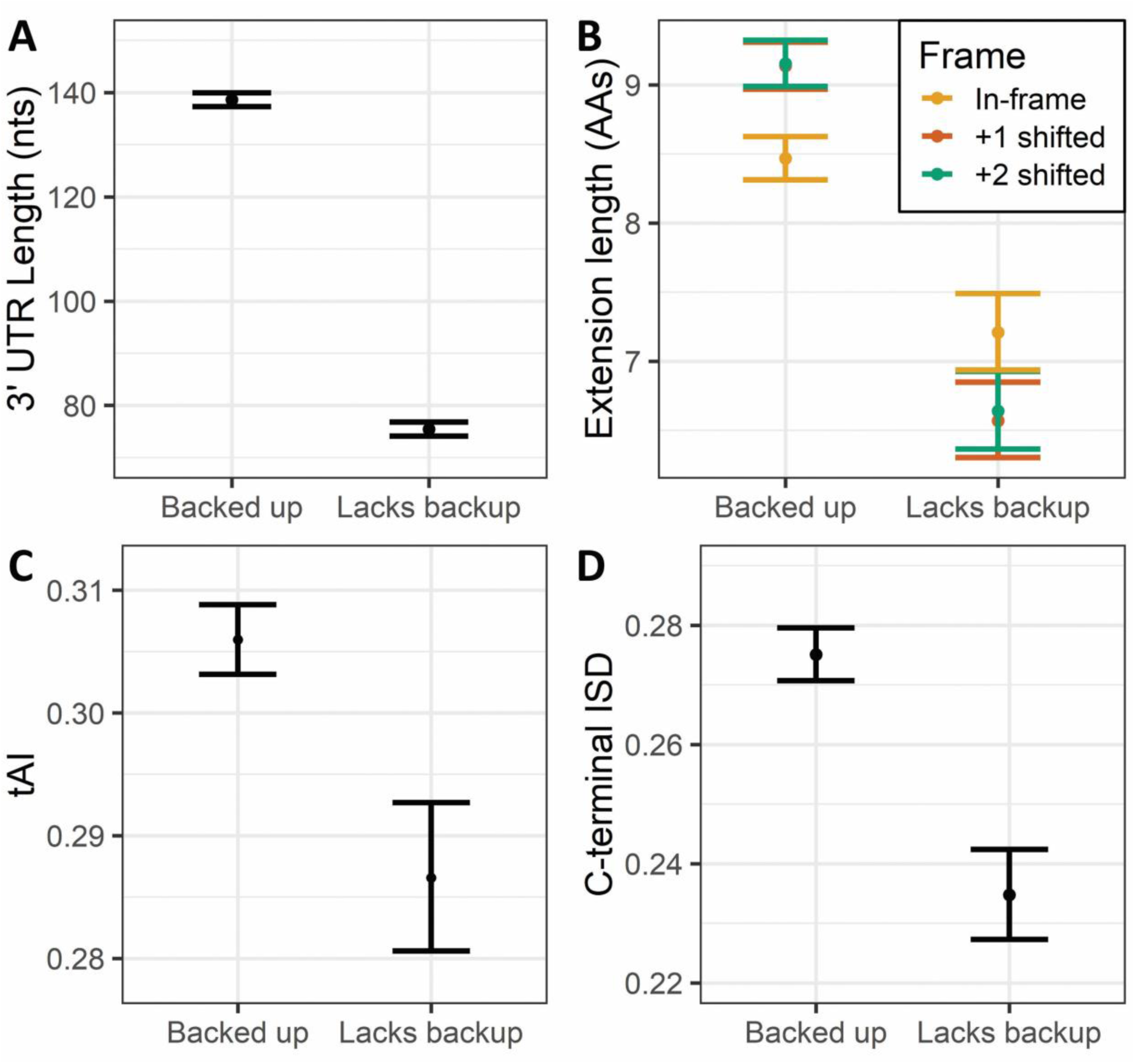
Proteins lacking a backup stop codon in at least one frame have shorter 3′ UTRs and extensions, more ordered C-termini, and slower in-frame codons beyond the stop codon. A) Proteins lacking backup(s) have shorter 3′ UTRs (P = 8 × 10^-191^, Student’s t-test on log-transformed 3′ UTR length). B) Proteins lacking backup(s) have shorter extension(s) in those frame(s) for which they do have a backup stop codon (P = 3 × 10^-4^, 1 × 10^-12^, and 1 × 10^-13^ for in-frame, +1 frame, and +2 frame extensions, respectively, Student’s t-test on log-transformed extension length). In-frame extensions are also shorter than both +1 shifted and +2 shifted extensions to proteins with fully backed up stop codons (P = 0.004 and 0.003, respectively, Student’s t-test on log-transformed extension lengths), but extension lengths between frames are not significantly different in proteins lacking backup(s) (P > 0.1 for both). C) The first four codon positions after the stop codon create slower in-frame translation in proteins that lack backup(s) than in proteins with fully backed up stop codons (P = 0.005, Student’s t-test on geometric mean tAI). D) Proteins lacking backup(s) have more ordered C-termini, assessed as the last 10 amino acids of the ORF (P = 5 × 10^-6^, Student’s t-test on square-root-transformed ISD). Error bars represent +/- one standard error. 3′ UTRs, extension lengths, and codon tAI values were log-transformed, and ISD was root-transformed, to calculate means and standard errors, and then back-transformed for the figure.

Genes that lack a backup in one frame are more likely to lack a backup in another, even after controlling for 3’UTR length and nucleotide content. To show this, we shuffled the nucleotides of each 3’UTR that was missing a backup stop codon in frame Y, and across 100,000 shufflings, recorded the simulated probability of lacking a backup in frame X, conditional on lacking it in frame Y. Then we generated a distribution of how many of these genes were expected to lack a backup in frame X, by summing over a Bernoulli sample from each gene-specific probability, repeating 10,000 times. The real genes had fewer backups in frame X than the simulated versions, for all frames X and Y, with p-values ranging from <10^-4^ to 0.01.

If this lack of readthrough is due to non-stop decay, we expect genes lacking a backup stop codon to be enriched for activity of Dom34, which is involved in ribosome rescue and is thought to be important for non-stop decay (Tsuboi, et al. 2012). Dom34 target genes, which exhibit elevated 3′ UTR ribohits in a Dom34 knockout strain, were identified by Guydosh and Green (2014) in a dataset independent from ours. Genes lacking at least one backup stop codon in our dataset are indeed enriched for Dom34 target genes relative to our full group of proteins (Table 1, P = 6.3 × 10^-47^, Pearson’s Chi-squared test on contingency table). Note that Dom34 target genes were only about 7-fold less likely than other genes to have a 3′ UTR ribohit in our dataset; lacking a backup stop codon is more predictive.

**Table 1.** Genes lacking a backup stop codon in at least one frame are enriched for Dom34 targets (P = 6.3 × 10^-47^, Chi-squared test).

| GENES WHERE THE 3' UTR<br>CONTAINS: | BACKUP STOP CODON (IN<br>ALL THREE FRAMES) | NO BACKUP STOP CODON (IN<br>AT LEAST ONE FRAME) |
| --- | --- | --- |
| KNOWN DOM34 TARGET | 54 | 122 |
| NOT A KNOWN DOM34 TARGET | 3055 | 851 |

When a protein lacking a backup stop codon in one frame is read through in a frame in which it does have a backup stop codon, other decay pathways could be employed. Decay might occur via no-go decay (Simms, et al. 2017) or slowness-mediated decay (Radhakrishnan, et al. 2016; Rak, et al. 2018) if inefficient codons in the extension cause slow or stalled translation, and hence a backup of ribosomes. We therefore hypothesize that proteins that lack backup stop codon(s) following frameshift employ slow codons immediately after the ORF stop codon. This prediction is restricted to in-frame extension codons because the location of the slow codons can be predicted, in contrast to frameshifts that can occur at many different positions prior to the stop codon. We assess codon speed using the tRNA adaptation index (tAI) (dos Reis, et al. 2004), a measure of codon preference based on tRNA copy number. With speed dominated by the slowest codon, we calculate geometric mean tAI (see Materials and Methods) for codon positions 1 to *n*. Proteins lacking out-of-frame backup(s) have more slowly translated in-frame extensions for values of *n* between 2 and 12 (P < 0.05 in each case, Student’s t-test). The effect size was largest for *n*=4, which we show in fig. 3C.

In the Introduction, we hypothesized that when the C-terminus is highly structured, extensions are likely to be disruptive. In agreement with this, ISD is lower in the last 10 amino acids of the ORFs of proteins lacking backup(s) than of fully backed up proteins (fig. 3D). We found no significant Gene Ontology enrichments (Ashburner, et al. 2000; Carbon, et al. 2017; Mi, et al. 2017) among proteins lacking backup(s) compared to our full set of analyzed proteins (P > 0.05, Fisher’s exact test with Benjamini-Hochberg false discovery rate correction).

Our central interest in this paper is in pre-adapting selection on sequences beyond stop codons. This interest is motivated by pre-adapting selection’s potential to contribute to the *de novo* evolution of protein-coding sequences. Fragile proteins are unlikely to ever contribute, and so are excluded, as identified by the absence of a backup stop codon, from the following analyses. So far we have demonstrated that selection has minimized the damage from readthrough of potentially fragile proteins. This selection on the readthrough of fragile proteins might be stronger than selection on non-fragile proteins, but it is the latter that we care about because of their potential for *de novo* evolution.

Similarly, we care most about the consequences of readthrough in the frame in which readthrough is most frequent. In-frame readthrough errors, driven by near-cognate tRNA pairings with stop codons, seem to occur more frequently than frameshift-driven readthrough errors. Specifically, work on nonsense suppression in reporter genes in *Escherichia coli* and other bacteria (Parker 1989), yeast (Namy, et al. 2001; Williams, et al. 2004), and mammals (Floquet, et al. 2012) has estimated wild-type in-frame readthrough rates of approximately 10^-2^ to 10^-4^. In contrast, stop-codon-bypassing +1 frameshifts occur on the order of 10^-4^ per translation event in *E. coli*, while +2 events occur on the order of 10^-5^ (Curran and Yarus 1986), and these observed frameshifting rates are likely elevated due to ribosome stalling associated with the premature nature of the stop codon. Stronger selection on the consequences of in-frame readthrough is supported in our data by the fact that in-frame extensions are shorter than both types of frameshift extension in proteins with fully backed up stop codons (fig. 3B). We therefore focus on in-frame readthrough of proteins with fully backed-up stop codons, and present these figures and tables in the main text. We also check extensions resulting from +1/+2 frameshifts at the stop codon, and present these figures and tables in our supplement.

### High ISD is a preadaptation

We primarily assess disordered status using IUPred2 (Dosztányi, et al. 2005; Meszaros, et al. 2018), except where noted. This program uses a sliding window to assess each amino acid’s interactions with its neighbors. We also sometimes assess or review other analyses of disorder at the level of individual amino acids in isolation, in terms of disorder propensity (Theillet, et al. 2013), relative solvent accessibility (RSA) (Tien, et al. 2013), or simply proportion of hydrophobic amino acids (where e.g. G, A, V, I, L, M, F, Y, and W are considered hydrophobic (Li and Zhang 2019)).

If high ISD is a preadaptation for C-terminal extensions, we expect coding sequences to display higher ISD than non-coding sequences. This prediction is supported for non-coding sequences beyond the stop codon (fig. 4A; previously found by Kleppe and Bornberg-Bauer (2018) for a high-readthrough subset of non-coding sequences, and by Li and Zhang (2019) for all genes using the proportion of hydrophobic amino acids), just as it was previously supported for mouse non-coding intergenic sequences (Wilson, et al. 2017) and alternative reading frames of viral coding sequences (Willis and Masel 2018). High ISD appears to be particularly important for C-termini, as disordered regions are most commonly found in the protein C-terminus (Uversky 2013), and adding hydrophobic amino acids to a C-terminus tends to lead to protein degradation (Arribere, et al. 2016). This is evidence that high ISD is indeed a preadaptation for C-terminal extensions.

**Fig. 4.**
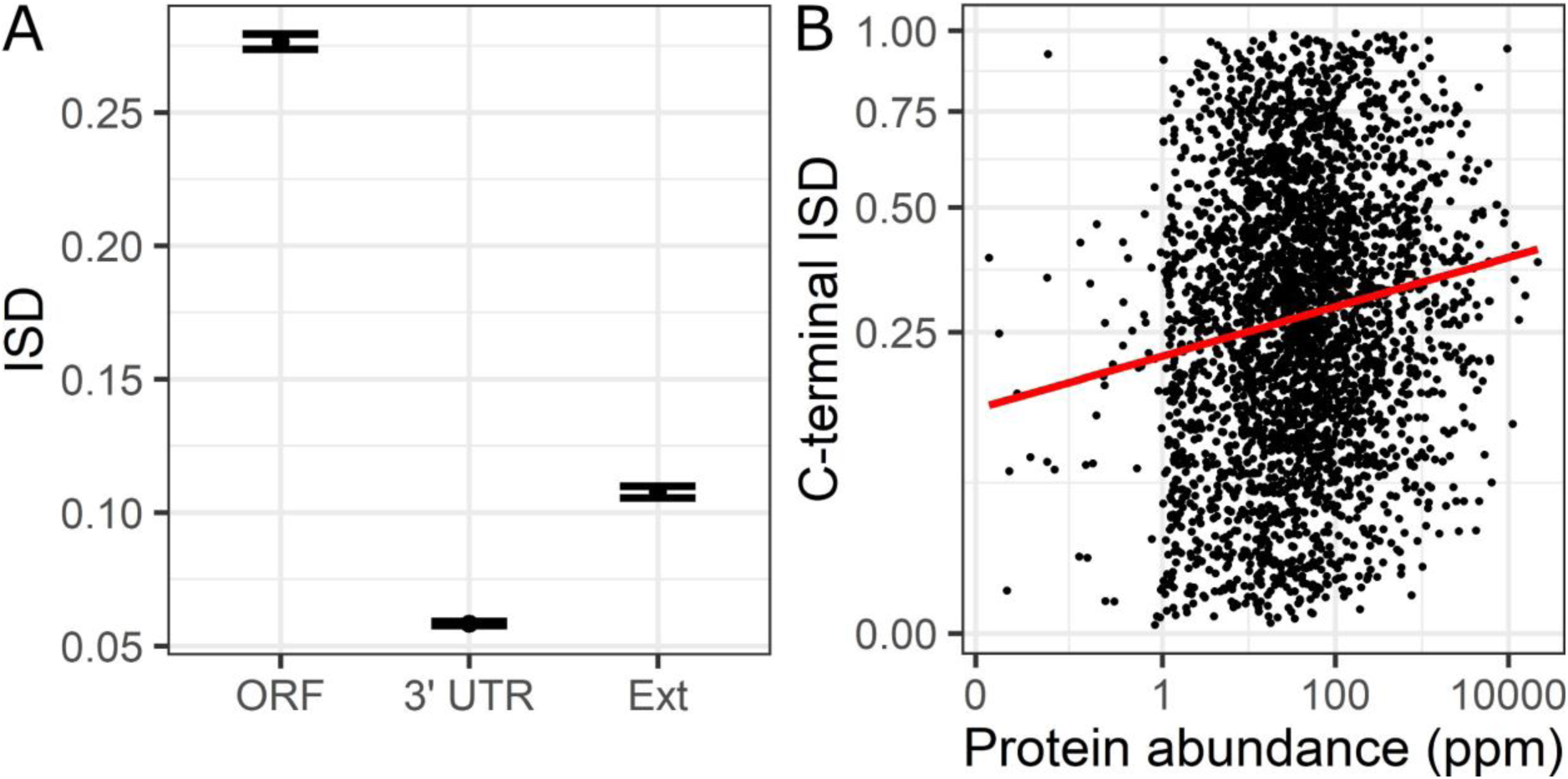
High ISD is a preadaptation. A) ISD is higher in coding regions than in non-coding sequences beyond the stop codon. Higher ISD in extensions than in complete 3′ UTRs is partly an artifact of the IUPred algorithm, which uses a sliding window in which C-terminal amino acids can affect the assessed ISD of extensions. Error bars represent +/- one standard error. ISD was root-transformed to calculate weighted means and standard errors, weighted by the length of the sequences, and then back transformed for the figure. B) More abundant non-fragile proteins have higher ISD in their C-termini. The regression line comes from a model where the root transform of the ISD of the last 10 amino acids of the ORF is predicted by log(protein abundance) (P = 9 × 10^-17^, likelihood ratio test). Note that while the data in B is very noisy (R^2^ = 0.022), the effect size is substantial, with a C-terminal ISD changing from 0.25 to 0.30 from the 25% to the 75% quantiles of protein abundance.

### Genes with detectable readthrough have higher ISD

Pre-adapting selection predicts that extensions that are read through more often will be more preadapted. In agreement with this prediction, in-frame extensions with more 3′ UTR ribohits have higher ISD (fig. 5A, Original), as do +2 shifted extensions to a slightly lesser degree (supplemental fig. 2A, Original) but not +1 shifted extensions (supplemental fig. 3A, see Discussion for an explanation as to why +1 frameshifts might be more naturally preadapted and have less need for pre-adapting selection). Note that Li and Zhang (2019) did not find low hydrophobicity in a set of 172 yeast genes suspected of having programmed readthrough. Our ribohit detection is more sensitive, i.e. we detect ribohits for many genes that Li and Zhang (2019) classed as non-readthrough, which might help increase our power.

**Fig. 5.**
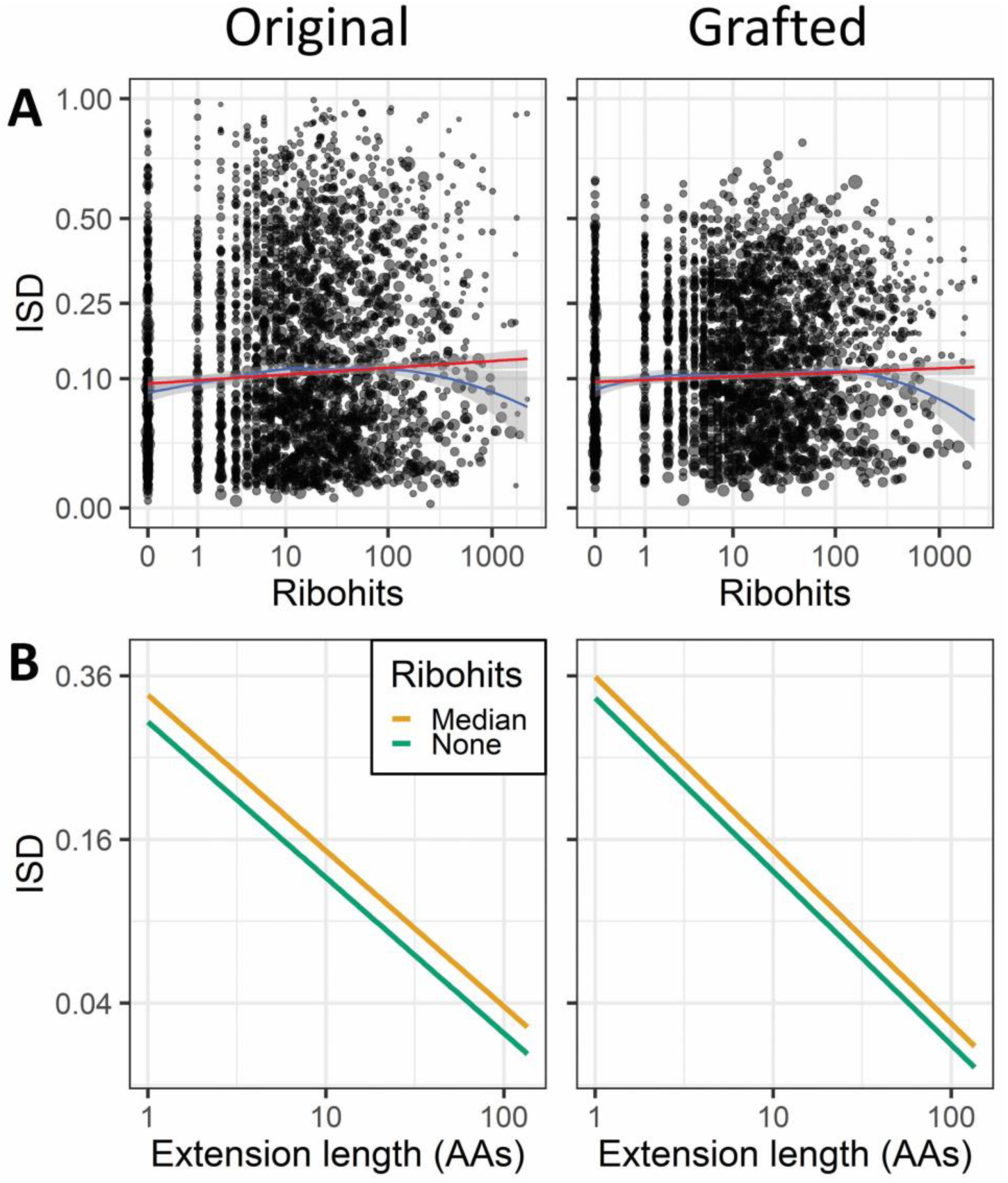
In-frame extensions with more 3′ UTR ribohits have higher ISD. A) Linear (red) and loess (blue) regressions of square-root transformed extension ISD on log-ribohits (without controlling for length). All regressions are weighted by the length of the extensions, visually represented by point area. The fact that after grafting (right) the curve persists with a similar slope, about 60% as steep, shows that elevated ISD in genes with more readthrough is at least partly driven by amino acids beyond the stop codon. Note that the downturn in the loess curve past 100 ribohits has wide confidence intervals (shaded gray), indicating sparse data, and is due to a few long length (and therefore high weight), low ISD extensions. In the non-weighted version of this curve (supplementary fig. 4A), the downturn is minimal, i.e. it is likely the result of an overemphasis on a few very long extensions that have many ribohits. Weighted R^2^ = 0.0036 and 0.0019 (Willett and Singer 1988), and R^2^ in the unweighted analysis = 0.0065 and 0.0027, for original and grafted, respectively. B) Ribohits still predict higher ISD (P = 4 × 10^-6^ and 3 × 10^-5^ for original and grafted extensions, respectively, in-frame ISD models, Table 2) after controlling for the large effect of extension length (P = 8 × 10^-50^ and 5 × 10^-90^). “Median” is the line for either the original (left) or grafted (right) in-frame ISD model in Table 2 when there are 13 3′ UTR ribohits, which is the median value for fully backed up proteins (see fig. 2), and “None” is for the same model with zero ribohits. Weighting by the length of the extensions (see Materials and Methods) means that the ISD values represent expectations from sampling an amino acid from the extensions rather than from sampling an extension.

ISD is driven in part by the interactions amino acids form with their neighbors (Dosztányi, et al. 2005); selection to differentially elevate ISD in protein C-termini could thus contribute to differentially elevated ISD in extensions, giving a false impression of pre-adapting selection on the extensions themselves. Kleppe and Bornberg-Bauer (2018) found that yeast proteins with more readthrough have more disordered C-termini. However, this relationship is at least partly driven by the low-readthrough, low-disorder fragile proteins described above, which do not contribute to protein innovation; once we exclude proteins lacking backup stop codons, genes with detectable readthrough still have higher ISD in the last 10 amino acids of the ORF, but this relationship is not statistically significant (P = 0.09, 2-tailed Student’s t-test on square-root transformed ISD). More convincingly, abundant proteins have higher ISD in their C-termini (fig. 4B, P = 9 × 10^-17^, see figure legend). This may be because selection shapes the C-terminus to mitigate the effects of readthrough, as suggested previously (Kleppe and Bornberg-Bauer 2018), or disordered C-termini may be intrinsically beneficial even in the absence of readthrough. This elevated protein ISD appears to be specific to the C-terminus; in a linear model where log abundance is predicted by the root transform of the ISD of the last 10 amino acids of the ORF, the root transformed ISD of the full ORF is no longer a significant predictor of protein abundance (P = 0.4, likelihood ratio test).

In order to test for pre-adapting selection on the extension sequences themselves, the effects of the C-terminal amino acids on the ISD of the extensions need to be controlled for. To do this, we grafted each extension onto the C-terminus of the same forty randomly selected non-fragile proteins, and took the average transformed ISD of the extension across those 40 standardized contexts. The effects of the extension sequences themselves are responsible for ≈60% of the readthrough-associated elevation in ISD in-frame (fig. 5A, Grafted has ≈60% the effect size of Original) and 50% of the elevation in ISD for +2 shifted extensions (supplemental fig. 2A).

IUPred accounts for nearby amino acid interactions via a 21 amino acid sliding window (Dosztányi, et al. 2005; Meszaros, et al. 2018), meaning that the first 10 amino acids of an extension are influenced by the last 10 amino acids on the ORF. Because proteins have higher ISD than expected from non-coding sequences, shorter extensions have more influence from the ORF and thus higher ISD by reason of short length alone. Thus, the results in fig. 5A and supplemental fig. 2A could be driven by extension length rather than ISD.

The number of ribohits is also confounded by extension length: there are more opportunities to detect ribohits when extensions are longer. Because high ISD is associated with short extensions which are associated with fewer ribohits, this negative confounding relationship makes our attempts to detect pre-adapting selection on ISD (i.e. higher ISD with more ribohits) conservative. However, selection on short extension length might create a positive confounding relationship between ribohits and ISD (mediated by short extensions), and this might swamp the negative confounding relationship between ribohits and ISD already described. This could lead to high ISD with more ribohits even in the absence of pre-adapting selection on ISD. In this case, we may mistake selection for short length for pre-adapting selection on ISD.

We therefore control for the effects of extension length on ISD as part of a linear regression model, as described in “ISD model” in the Methods. Extensions with more 3′ UTR ribohits have higher ISD even after controlling for length for both in-frame (fig. 5B; P = 3 × 10^-5^, grafted in-frame ISD model, Table 2) and +2 shifted extensions (supplemental fig. 2B, P = 5 × 10^-4^, grafted +2 frame ISD model, supplemental Table 1). Controlling for length strengthened rather than weakened our results: including the effects of C-termini, the length-controlled slope on ribohits is about 1.4 and 1.3 times the non-length-controlled slope for in-frame and +2 shifted extensions, respectively, and about 1.7 times after grafting for both in-frame and +2 shifted extensions.

**Table 2.**
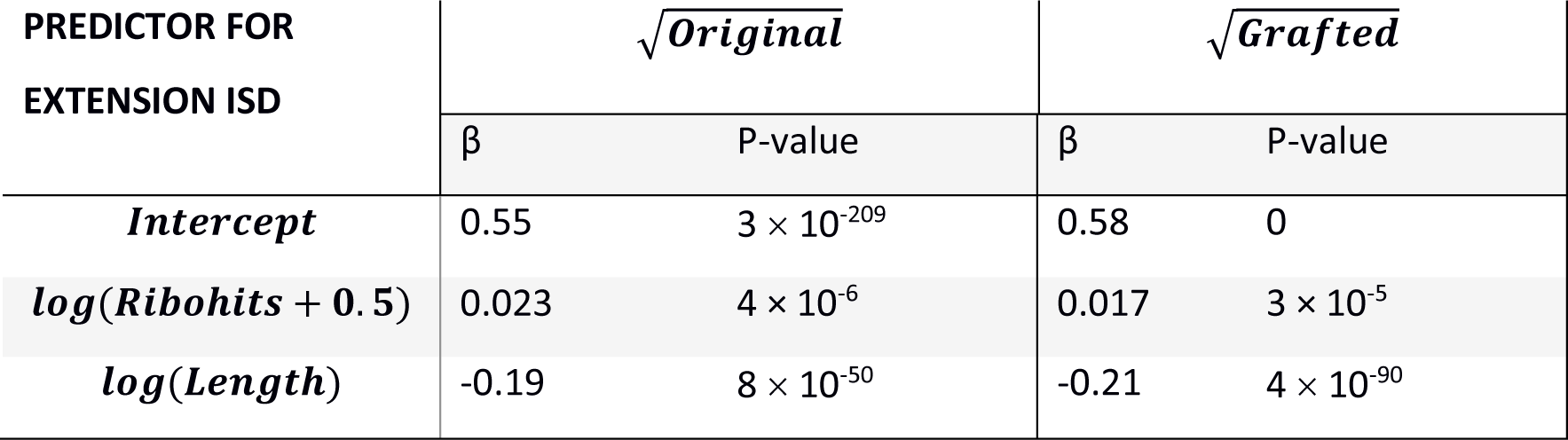
Regression model predictors of in-frame original and grafted ISD

| PREDICTOR FOR<br>EXTENSION ISD | $\sqrt{\text{Original}}$ | | $\sqrt{\text{Grafted}}$ | |
| --- | --- | --- | --- | --- |
| | $\beta$ | P-value | $\beta$ | P-value |
| *Intercept* | 0.55 | $3 \times 10^{-209}$ | 0.58 | 0 |
| *log(Ribohits + 0.5)* | 0.023 | $4 \times 10^{-6}$ | 0.017 | $3 \times 10^{-5}$ |
| *log(Length)* | -0.19 | $8 \times 10^{-50}$ | -0.21 | $4 \times 10^{-90}$ |

### Amino acids after stop codon are shaped by pre-adapting selection

The ISD results above involve complicated methods to control for confounding factors, in particular extension length. Because IUPred uses a sliding window, the degree to which C-terminal amino acids elevate extension ISD depends on extension length, even after grafting. We therefore also use two straightforward amino acid scores that do not involve a sliding window: RSA, which scores how often amino acids are found in the interior vs. surface of globular proteins (Tien, et al. 2013), and disorder propensity (Theillet, et al. 2013), which scores how often amino acids tend to be found in disordered regions compared to ordered regions. As predicted by pre-adapting selection, in-frame extensions with more ribohits have higher RSA (fig. 6A, left) and disorder propensity (fig. 6A, right), as do +2 shifted extensions (supplemental fig. 5A) but not +1 shifted extensions (supplemental fig. 6A). Surprisingly, and in contrast to results for extension disorder as a whole, the +2 effect sizes are slightly larger than in-frame: about 1.05 times larger for RSA and 1.4 times larger for disorder propensity.

**Fig. 6.**
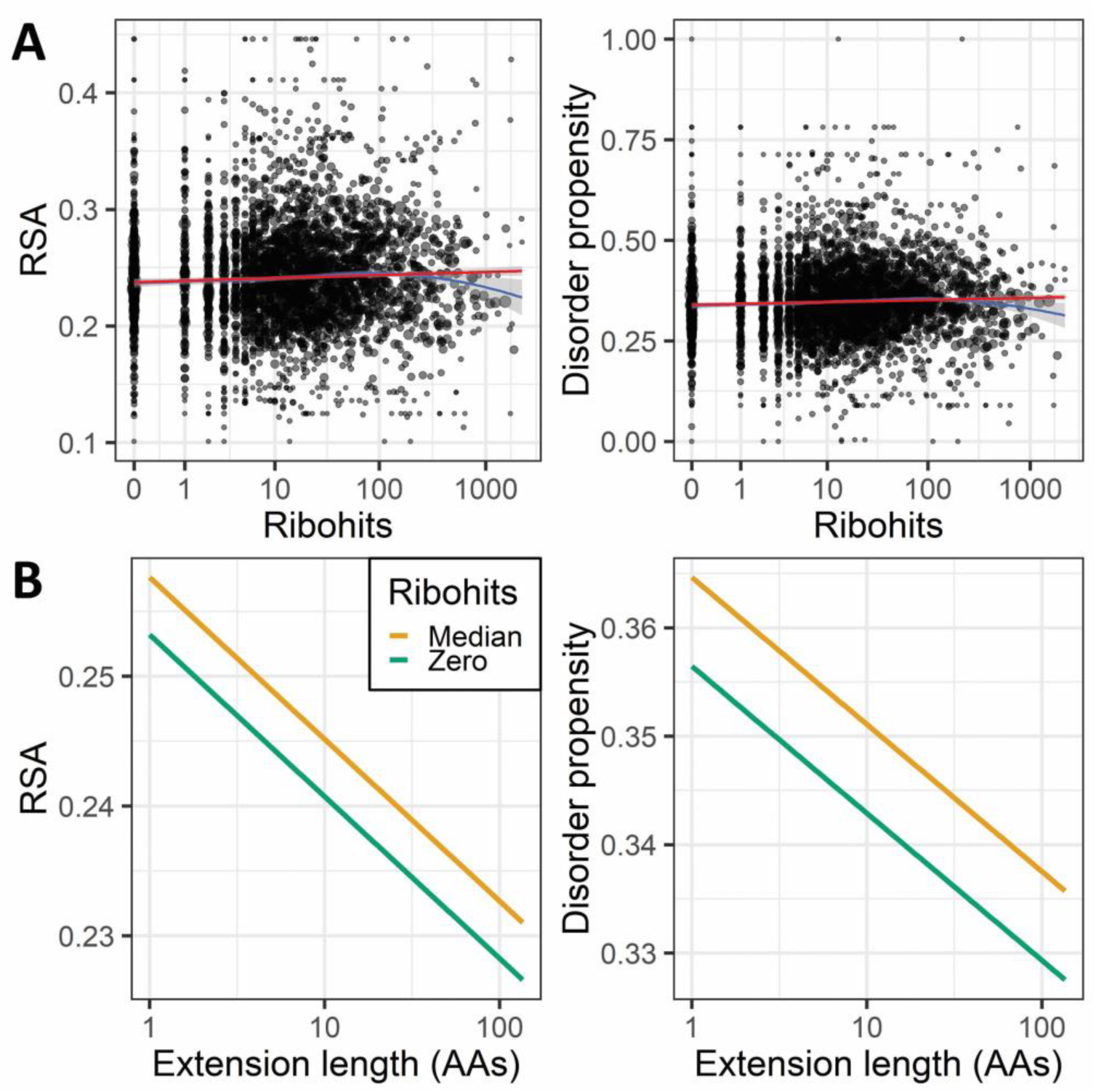
Metrics that do not use a sliding window provide further evidence that pre-adapting selection acts on extension amino acids. A) Linear (red) and loess (blue) regressions of in-frame mean RSA (left) and disorder propensity (right) on log-ribohits (without controlling for length). All regressions are weighted by the length of the extensions, visually represented by point area. In the non-weighted version of this curve (supplementary fig. 7A), the downturn between 100 and 1000 ribohits is minimal, i.e. it is likely the result of an overemphasis on a few very long extensions that have many ribohits. Weighted RM^2^ = 0.0028 and 0.0026 (Willett and Singer 1988), and R^2^ in the unweighted analysis = 0.0013 and 0.00073, for RSA and disorder propensity, respectively. B) After controll controlling for the effect of extension length, mean in-frame extension RSA and disorder propensity are still higher with more 3′ UTR ribohits (P = 5 × 10^-4^ and 0.002, RSA and disorder propensity models, respectively, Table 3). Extension length is also predictive; mean extension RSA and disorder propensity are both lower for longer extensions (P = 3 × 10^-8^ and 0.003). For a model with both extension length and ribohits as predictors of RSA and disorder propensity, respectively, weighted R^2^ = 0.013 and 0.0052 (Willett and Singer 1988), and in the corresponding unweighted analyses R^2^ = 0.012 and 0.0077. Lines for “Median” and “None” are as described in fig. 5, except using either the RSA (left) or disorder propensity (right) model in Table 3. As in fig. 5, weighting by the length of the extensions means that RSA and disorder propensity values represent expectations from sampling an amino acid from the extensions rather than from sampling an extension. Note that the effect size in B) here appears larger than the effect size in fig. 5B, but this is misleading; the effect size in 5B is actually larger but appears smaller because it is dwarfed by the large effect size of extension length on ISD.

Short in-frame extensions also have higher RSA (fig. 6B, left) and disorder propensity (fig. 6B, right), in agreement with suggestions that they are less likely to lead to protein degradation (Arribere, et al. 2016). The same is even more true for +2 shifted (supplemental fig. 5B; 1.7 times and 1.6 times the effect size as for in-frame RSA and disorder propensity, respectively) and +1 shifted extensions (supplemental fig. 6B; 3.6 times and 4.1 times the effect size as for in-frame RSA and disorder propensity, respectively). The effect of length on RSA and disorder propensity was notably larger than the effect of ribohits. Specifically, the change in in-frame extension RSA from the 25% to the 75% quantiles of length was 2.9 times as large as from the 25% to the 75% quantile of ribohits, and 1.7 times for disorder propensity; similar comparisons for +2 shifted extension RSA and disorder propensity yielded 4.6 and 2.0, respectively. Shortness of extension might thus be a better metric of the quantity of readthrough products subjected to pre-adapting selection than the number of ribohits.

## Discussion

The pre-adapting selection hypothesis predicts that the more often an erroneous gene product is produced, the more likely it is to be preadapted. We found that non-fragile proteins that are read through more often have C-terminal extensions with higher disorder. These high disorder extensions constitute a pool of preadapted cryptic sequences that evolution could draw from for *de novo* evolution, in a C-terminus extension process known to occur (Giacomelli, et al. 2007; Vakhrusheva, et al. 2011; Andreatta, et al. 2015)).

Higher readthrough could impose pre-adapting selection on a C-terminal extension, or a fortuitously preadapted C-terminal extension could alleviate selective pressure against readthrough – our methods are not able to conclusively determine the causal direction. However, we have two pieces of evidence to suggest that selection is operating in the direction posited by pre-adapting selection. First, in-frame extensions in genes with fully backed up stop codons are shorter than frameshifted extensions (fig. 3B). A few fortuitously short extensions are unlikely to change the fact that most readthrough occurs in-frame, so the fact that in-frame extensions tend to be shorter than frameshifted extensions (fig. 3B) in proteins with fully backed up stop codons indicates causality in the appropriate direction. Second, observed readthrough reflects a combination of leakiness (probability of readthrough per translation event), protein abundance, and degradation of readthrough products. The consequences of readthrough are unlikely to be the most significant factor shaping the evolution of protein abundance, and so the problematic scenario, in which causation works in the opposite direction to that posited by pre-adapting selection, is presumably restricted to selection on leakiness and degradation. We found that more abundant proteins tend to have shorter in-frame extensions (P = 4 × 10^-6^, Pearson’s correlation coefficient = −0.083), further suggesting that a causal direction from readthrough to selection on extensions is valid, at least for in-frame extensions.

Evidence for pre-adapting selection based on extension ISD was, as expected, stronger for in-frame extensions than for frameshifted extensions. Consistent with this is the high frequency with which in-frame extensions are incorporated into C-termini during evolution (Giacomelli, et al. 2007). Part of this stronger in-frame effect comes from the C-terminal itself, whose identity is less consistent for frameshifts given variation in frameshift location. Perhaps for this reason, pre-adapting selection on the RSA and disorder propensity of amino acids beyond the stop codon, which excludes any influence of the C-terminal itself, is slightly stronger +2 than in-frame.

Surprisingly, we found evidence for pre-adapting selection on +2 shifted extensions but not on +1 shifted extensions, despite other evidence that the latter form of readthrough occurs at higher rates (reviewed in Results). A possible explanation is that a bias towards hydrophilicity makes the +1 shifted extensions more naturally preadapted, alleviating the need for pre-adapting selection, while +2 frame extensions are biased toward hydrophobicity. The bias stems from the identity of the codons created by a frameshifted stop codon. If a +1 frameshift occurs upstream of the stop codon, the T beginning the stop codon becomes the last nucleotide of the first codon of the extension, introducing little bias. But whether the +1 frameshift occurs at or prior to the stop codon, it creates a codon that begins with AA, AG, or GA, which creates a bias towards hydrophilicity. Specifically, AAY and AGY encode asparagine and serine, both of which are polar, while AAR, AGR, GAY, and GAR encode lysine, arginine, aspartic acid, and glutamic acid, respectively, all of which are charged. This guaranteed hydrophilicity increases the ISD of the +1 extension, alleviating the need for pre-adapting selection. For +2 frameshifts, one codon must end with TA or TG when the frameshift occurs upstream of the stop codon, while the next codon must start with A or G. While the latter introduces little bias, codons ending in TA code for leucine, isoleucine, and valine, while codons ending in TG code for leucine, methionine, and valine. All of these are strongly hydrophobic, creating a bias toward low ISD in real extensions generated by upstream +2 frameshifts, albeit not captured by our ISD metrics based on frameshifts occurring precisely at the stop codon. The extra hydrophobicity of these +2 shifted extensions may make them less likely to be preadapted, exacerbating the need for pre-adapting selection. As for in-frame extensions, our ISD calculation simply excised stop codons to generate in-frame extensions, but real extensions usually result from a near-cognate tRNA pairing with a stop codon. The most common near-cognate decodings in yeast are tyrosine or glutamine for UAA, tyrosine for UAG, and tryptophan for UGA (Blanchet, et al. 2014). Glutamine is hydrophilic, tyrosine is amphipathic, and tryptophan is sometimes grouped as hydrophobic and sometimes as amphipathic, collectively suggesting that bias will not be strong.

We found that extensions with more ribohits tend to be shorter, implying selective pressure for shorter extensions in budding yeast. Previous studies have also shown evidence of selection for shorter extensions in eukaryotes (but not in bacteria; Ho and Hurst 2019). In yeast, backup stop codons are more frequent than expected by chance alone (Williams, et al. 2004), show evidence of conservation (Liang, et al. 2005), and are overrepresented in genes with a high codon adaptation index (Liang, et al. 2005). However, shorter extensions are unlikely to contribute appreciably to *de novo* innovations, precisely because of their short length. We have not only confirmed that selection prefers short extensions in yeast, but also found for the first time that selection favors high ISD extensions. High ISD increases the potential for *de novo* innovation (Wilson, et al. 2017; Willis and Masel 2018).

Error rates are not constant throughout the genome. We found that fragile proteins have dramatically lower readthrough than non-fragile proteins. Lack of backup stop codons triggering non-stop mediated mRNA decay (Vasudevan, et al. 2002) due to the absence of a backup stop codon, and slow codons in extensions triggering no-go (Simms, et al. 2017) or slowness-mediated decay (Radhakrishnan, et al. 2016; Rak, et al. 2018), are two of the mechanisms by which this might occur. Another possible mechanism is an mRNA structure that physically blocks ribosome progression into the 3′ UTR, which would stall ribosomes and also lead to no-go decay (Harigaya and Parker 2010).

Many molecular errors are less common in highly expressed genes, including errors in transcription start site in human and mouse (Xu, et al. 2019), mRNA polyadenylation in mammals (Xu and Zhang 2018), post-transcriptional modifications in human and yeast (Liu and Zhang 2018b, a), and mistranscription errors in *E. coli* (Meer, et al. 2019). Most importantly for our study, readthrough errors in yeast are less common in highly expressed genes (Li and Zhang 2019). However, this does not remove selection for benign extensions that are more disordered.

Here we have provided a proof of principle that pre-adapting selection shapes the sequences beyond stop codons, facilitating later *de novo* evolution of C-termini. This makes it plausible that pre-adapting selection also works on the precursors of full *de novo* genes, junk polypeptides (Wilson and Masel 2011; Ruiz-Orera, et al. 2018; Blevins, et al. 2019; Durand, et al. 2019), facilitating later *de novo* birth of complete proteins. This can reconcile the perceived implausibility of *de novo* gene birth (Zuckerkandl 1975; Jacob 1977) with recent convincing evidence that *de novo* gene birth is a real phenomenon.

## Materials and Methods

### Data

Pre-processed ribosome profiling data on wild-type [psi^-^] and [PSI^+^] strains of *S. cerevisiae* was taken from Jarosz et al. (In prep), and is available at https://doi.org/10.6084/m9.figshare.11418789.v1. Jarosz et al. (In prep) used flash freezing to inhibit elongation according to a modified protocol of Brar et al. (2012).

We chose Jarosz et al.’s (In prep) dataset for four reasons: 1) it has very high coverage with a substantial number of reads mapping to 3′ UTR regions (fig. 2; overall coverage is higher than (e.g. Baudin-Baillieu, et al. 2014; Cheng, et al. 2018; Spealman, et al. 2018)), 2) it is methodologically designed to minimize spurious 3′ UTR ribohits, such as avoiding use of micrococcal nuclease which is known to elevate 3′ UTR ribohits (Miettinen and Björklund 2015), 3) it uses wild-type yeast rather than mutants that perturb translation termination (e.g. Nedialkova and Leidel 2015; Guydosh and Green 2017), and 4) it uses both [psi^-^] and [PSI^+^] strains. [PSI^+^] sequesters the translation release factor Sup35p, increasing the rate of stop codon readthrough (Cox 1965; Liebman and Sherman 1979; Firoozan, et al. 1991). This elevated readthrough is thought to be favored by selection for its role in evolvability (Masel and Bergman 2003; Lancaster, et al. 2010), making it biologically relevant. Conveniently, [PSI^+^] increases the rate of readthrough (Cox 1965; Liebman and Sherman 1979; Firoozan, et al. 1991) and hence our ribohit coverage in extensions across the board rather than in a gene-specific manner (Jarosz, et al. In prep). Note that Kleppe and Bornberg-Bauer (2018) used the data of Nedialkova and Leidel (2015), which was high coverage but primarily composed of mutants prone to ribosomal pausing due to wobble base-pairing deficiencies, so the wild type coverage was lower than the data used here. Li and Zhang (2019) used the data from Dunn et al. (2013), which was primarily focused on *Drosophila melanogaster* and appeared to have limited publically available yeast data.

Sequences for 6,752 protein-coding genes were downloaded from the Saccharomyces Genome Database (SGD, http://www.yeastgenome.org) using the 2011 S288C reference *S. cerevisiae* genome sequence and associated annotations, matching the strain used for ribosomal profiling and the gene annotations used to analyze that data (Jarosz, et al. In prep). 3′ UTR sequence annotations were done by Jarosz et al. (In prep) by taking the average 3′ UTR length from transcript isoform profiling data (Pelechano, Wei, and Steinmetz 2013). We excluded 2,032 genes without a 3′ UTR annotation, and a further 646 whose ORF was not “verified” as protein-coding in SGD, resulting in 4,083 genes with verified ORFs. Lastly, one gene, YOR031W, has a blocked reading frame with an in-frame stop codon after only eight amino acids; excluding it left 4,082 genes.

We scored how often a ribohit was found within the 3′ UTR region, including both [psi^-^] and [PSI^+^] strains. For our statistical analyses, we log transformed total 3′ UTR ribohits for each extension, which achieved an approximately linear relationship with ISD (fig. 5A and supplemental figs. 2A and 3A), RSA (fig. 6A and supplementary figs. 5A and 6A), and disorder propensity (fig. 6A and supplementary figs. 5A and 6A). Note that some of our genes had zero 3′ UTR ribohits – to include our zero 3′ UTR ribohits genes in our log transform, we added 0.5 to all genes’ 3′ UTR ribohits.

Protein abundance data was downloaded (11/30/15) from PaxDB (Wang, et al. 2012; Wang, et al. 2015) using the integrated *S. cerevisiae* dataset, expressed as parts per million individual proteins (ppm). Of the 4,082 genes in our analysis, abundance data was available for all but two; these two genes were excluded from analyses that used abundance data. Abundance values were log transformed before analysis; based on manual inspection, this achieved approximately normally distributed values.

In-frame readthrough occurs when a near-cognate tRNA pairs with a stop codon (Blanchet, et al. 2014). Rather than guess at tRNA pairings, we simply excised the stop codon and scored extensions as the codons between the stop codon and the next in-frame backup stop codon. When calculating in-frame extensions lengths, we included the unknown amino acid corresponding to the stop codon itself.

+1 and +2 frameshifted extensions were scored by shifting the reading frame downstream by one or two nucleotides, respectively, at the stop codon. +1 and +2 frameshifts therefore include either the last two or last one of the stop codon nucleotides, respectively, in the extension. Real frameshifts do not have to happen at the stop codon and can occur at any location along the ORF, leading to an extension rather than a truncation if no stop codon is created between the location of the frameshift and the position of the original stop codon. Our analysis was restricted to frameshifts that happen at the stop codon, both for convenience, and because these do not disrupt the normal amino acids of the ORF and are the most likely to be at least relatively benign. If at least one frame had no backup stop codon in the 3′ UTR as annotated by Jarosz et al. (In prep), that gene was placed in our “lacks backup” class of proteins (see figs. 2 and 3).

All in-house scripts for our analyses can be found at https://github.com/MaselLab/Kosinski-and-Masel-CTerminalExtensions. All figures were made in R (R Core Team 2019) using the “ggplot2” package (Wickham 2016).

### Scoring ISD

We used the “long” setting in IUPred2 (Dosztányi, et al. 2005; Meszaros, et al. 2018), which returns a score between zero and one for each amino acid in a sequence. These scores represent non-independent quasi-probabilities that the amino acid is in a disordered region. We estimate the ISD of a sequence using a simple average of the IUPred2 quasi-probabilities for the amino acids in the sequence of interest. Because this procedure creates heteroscedasticity, by making the error a function of sequence length, we use sequence length as a weight in later analyses. Note that the unknown near-cognate amino acid that pairs with the stop codon was not scored for its ISD, RSA or disorder propensity. Consequently, we did not count this unknown amino acid as part of in-frame extension length when using in-frame extension length as a weight for in-frame extension ISD/RSA/disorder propensity (but it was counted as part of in-frame extension length at all other times).

To calculate the ISD of the last 10 amino acids of an ORF, we fed the full ORF into IUPred2, then took the average only of the last 10 amino acids. Similarly, to calculate the ISD of extensions, we fed the sequence up to the backup stop codon into IUPred2, and took the average score only of the amino acids in the extension.

Square-root-transformed ISD values were approximately normal for ORF, C-terminal, and grafted extension ISD, and were used in subsequent statistical analyses. This corresponds to the biological intuition that an ISD difference of 0.7 vs 0.8 is less important than a difference between 0 and 0.1. Note that we transformed sequence ISD; we did not transform IUPred2 quasi-probabilities prior to averaging to produce sequence ISD.

IUPred2 was downloaded on April 7^th^, 2018.

### RSA calculation

Relative solvent accessibility (RSA) is a measurement of how often an amino acid tends to be found in the exterior versus interior of globular proteins (Tien, et al. 2013), calculated from high quality crystal structures using the DSSP program (Kabsch and Sander 1983; Tien, et al. 2013). More hydrophilic residues tend to be found on the exterior.

### tAI

tAI was calculated for each codon in a sequence using a script provided by Roni Rak, and tRNA copy numbers for *S. cerevisiae* were taken from Percudani et al. (1997). Standard practice for calculating sequence tAI is to use the geometric mean (dos Reis, et al. 2004). Equivalently, we calculated sequence tAI as the arithmetic mean of the log-transformed tAI codon scores.

### Regression models

During model building, we used log_10_ transforms of our predictors when this gave a better model fit. Tests of statistical significance for all models came from the likelihood ratio test associated with dropping the variable of interest from its respective regression model shown in Tables 2-3. P-values reported from linear models come from models controlling for all predictive factors listed below, not just the predictive factors mentioned in the Results until that point. All listed predictor coefficients (β) come from the model with all listed predictors included.

**Table 3.** Regression model predictors of in-frame mean RSA and disorder propensity.

| PREDICTOR FOR<br>RSA/DISORDER | RSA |  | Disorder propensity |  |
| --- | --- | --- | --- | --- |
| | $\beta$ | P-value | $\beta$ | P-value |
| *Intercept* | 0.25 | 0 | 0.36 | 0 |
| *log(Ribohits + 0.5)* | 0.0031 | $5 \times 10^{-4}$ | 0.0057 | 0.002 |
| *log(Length)* | -0.012 | $3 \times 10^{-8}$ | -0.0014 | 0.003 |

#### ISD models

A separate model was built to predict root-transformed ungrafted ISD values and for average root-transformed grafted ISD values (Table 2). ISD is a mean across extension amino acids, so we weight each value by the length of the extension. We tested the effects of log (ribohits + 0.5) while controlling for extension length. For +2 and +1 shifted extension ISD models, see supplementary tables 1 and 2, respectively.

#### RSA and disorder propensity models

We built models for extension mean RSA and for disorder propensity in the same way as for ISD (see section above), including weighting (Table 3). Manual inspection showed that both RSA and disorder propensity were approximately normal, and hence neither was transformed. For +2 and +1 shifted extension mean RSA and disorder propensity models, see supplementary tables 3 and 4, respectively.

## Acknowledgements

This work was supported by the John Templeton Foundation (39667, 60814) and the National Institutes of Health (GM-104040, T32GM-008659). We thank Dan Jarosz for sharing his data with us ahead of publication and for his insights concerning ribosome profiling, Adam Hancock, Jason Slepicka, and Taylor Kessinger for their work on an earlier version of this project fatally handicapped by a lack of such good data, Premal Shah for sharing pipelines that we didn’t end up using after getting Dan Jarosz’s data, Alex Lancaster for sharing his ribosome profiling scripts that we ended up not using, Roni Rak and Tzachi Pilpel for sharing their script and insights on tAI, the Stephen Floor lab for suggesting that we examine Dom34 target genes, and Eden Eaton and Scott Foy for contributions during the early stages of this project.

## Notes

#### Summary of Updates

Figure 2 moved from supplement to main paper to replace table 1. Updated figures 5A and 6A (along with similar supplemental figures) to have point area proportional to weight, and added supplemental figures to show unweighted regressions. Added R-squared values for weighted and unweighted regressions. Clarified our logic for figure 3. Made various minor edits to the text.

https://github.com/MaselLab/Kosinski-and-Masel-CTerminalExtensions

https://doi.org/10.6084/m9.figshare.11418789.v1

