## Supplemental materials for "Readthrough errors purge deleterious cryptic sequences, facilitating the birth of coding sequences"

### Supplemental figures

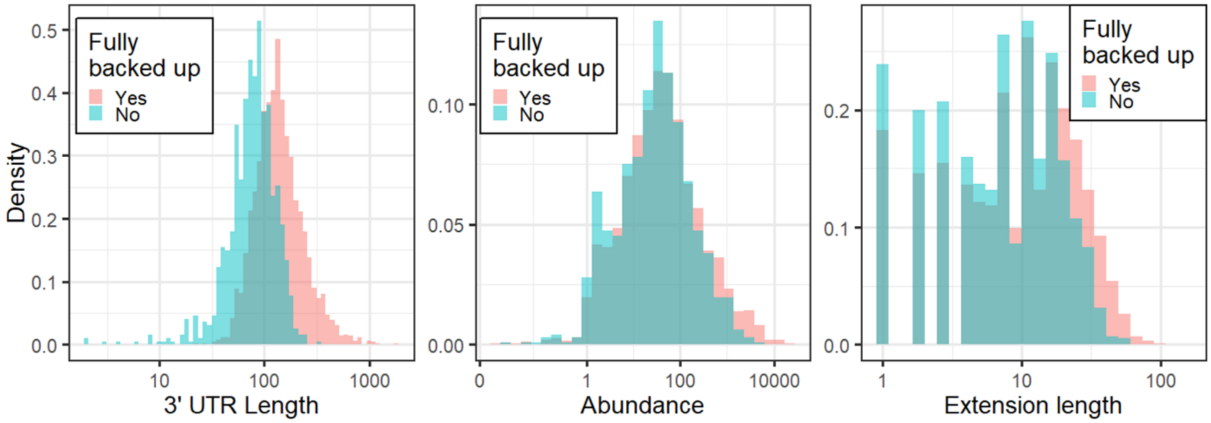


**Supplemental fig. 1.** Fully backed up and not fully backed up proteins exhibit substantial overlap in their distributions of 3′ UTR lengths, protein abundances, and extension lengths.

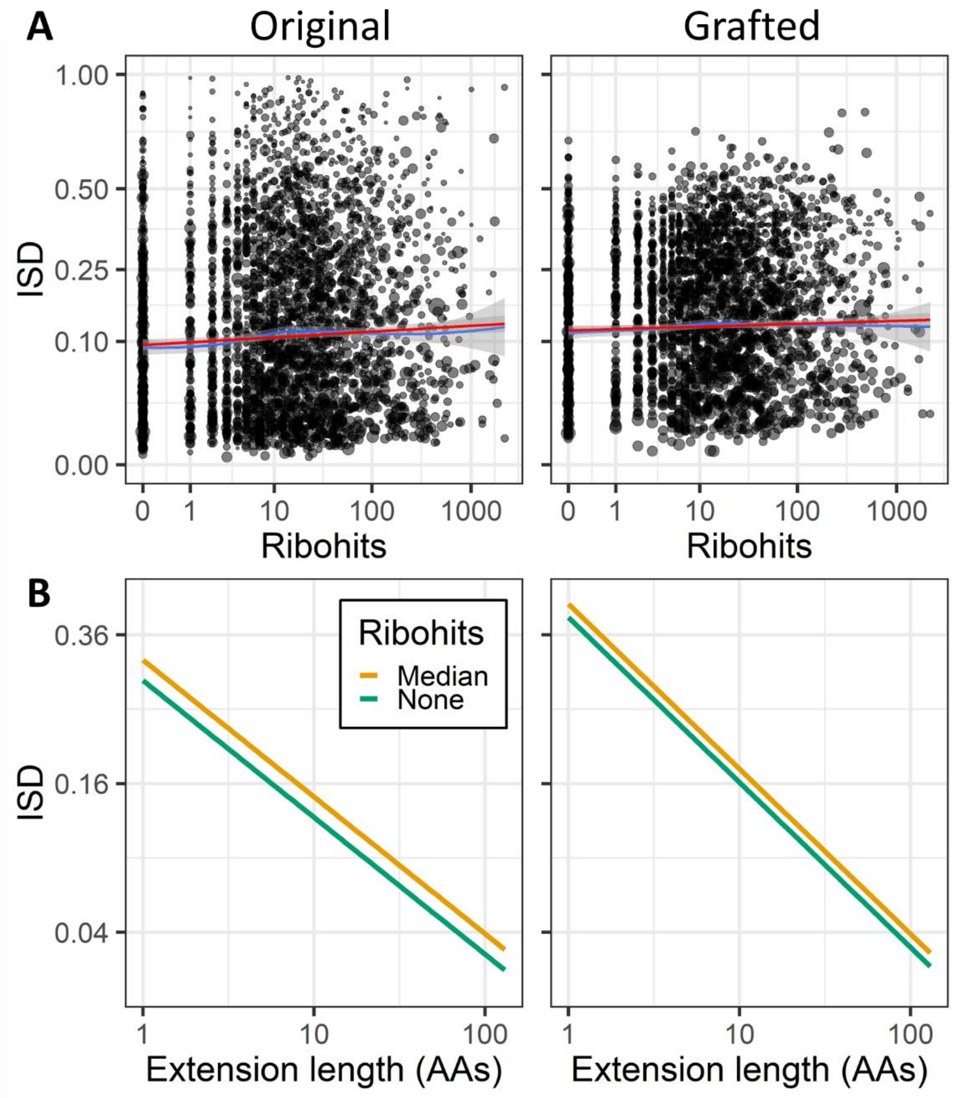


**Supplemental fig. 2.** +2 shifted extensions with more 3′ UTR ribohits have higher ISD. A) Linear (red) and loess (blue) regressions of square-root transformed extension ISD on log-ribohits (without controlling for length). All regressions are weighted by the length of the extensions, visually represented by point area. The fact that after grafting (right) the curve persists with a similar slope, about 50% as steep, shows that elevated ISD in genes with more readthrough is at least partly driven by amino acids beyond the stop codon. Weighted $R^2 = 0.0029$ and 0.00079 (Willett and Singer 1988), and in the corresponding unweighted analyses $R^2 = 0.0043$ and 0.00053, for original and grafted, respectively. B) Ribohits still predict higher ISD ($P = 2 \times 10^{-5}$ and $5 \times 10^{-4}$ for +2 shifted original and grafted extensions, respectively, +2 frame ISD models, supplemental Table 1) after controlling for the large effect of extension length ($P = 2 \times 10^{-57}$ and $6 \times 10^{-120}$). “Median” is the line for either the original (left) or grafted (right) +2 shifted ISD model in supplemental Table 1 when there are 13 3′ UTR ribohits, which is the median value for fully backed up proteins (see fig. 2), and “None” is for the same model with zero ribohits. Like in fig. 5, weighting by the length of the extensions (see Materials and Methods) means that the ISD values represent expectations from sampling an amino acid from the extensions rather than from sampling an extension.

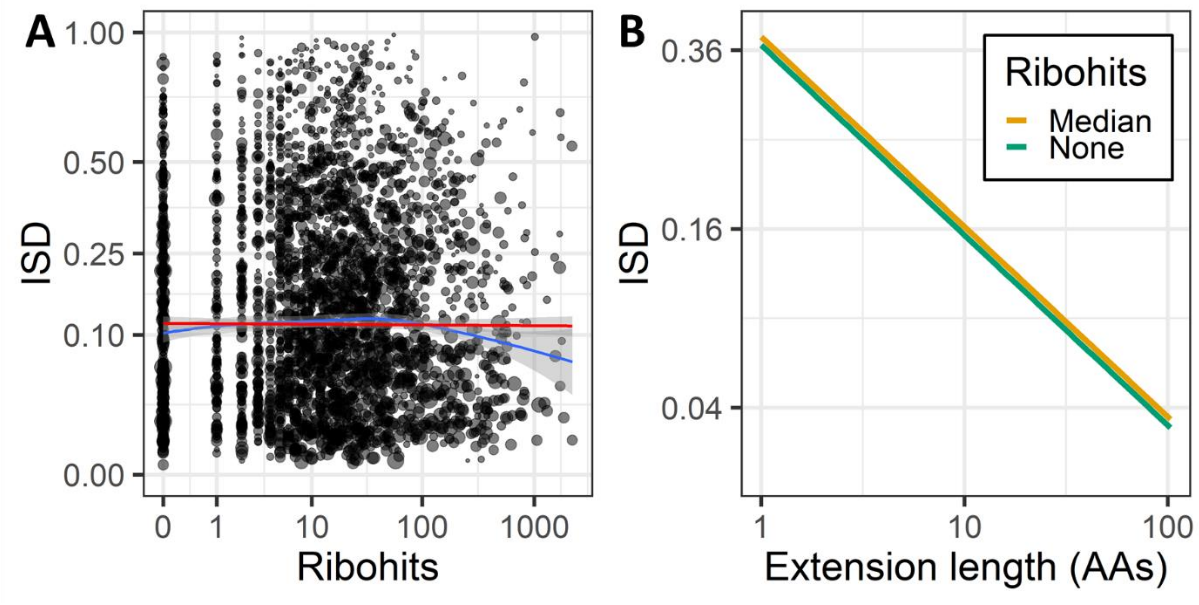


**Supplemental fig. 3.** +1 shifted extension ISD does not display evidence of pre-adapting selection. A) Linear (red) and loess (blue) regressions of square-root transformed extension ISD on log-ribohits without controlling for length are shown. Both regressions are weighted by the length of the extensions, visually represented by point area. B) After controlling for extension length, log ribohits are not a significant predictor of +1 extension ISD ($P > 0.1$, +1 shifted original extension ISD model, Table 2). Ribohits and extension length are log transformed, while +1 extension ISD is square-root transformed. Lines for "Median" and "None" are as described in supplemental figure 2, except using the +1 shifted extension ISD model in supplemental Table 2. Note that grafted extensions are not shown for +1 shifted extensions; because the non-grafted (i.e. original) +1 shifted extension ISD do not show a significant effect with ribohits, there was no need to show whether a non-significant effect persisted after controlling for the effects of C-terminal amino acids.

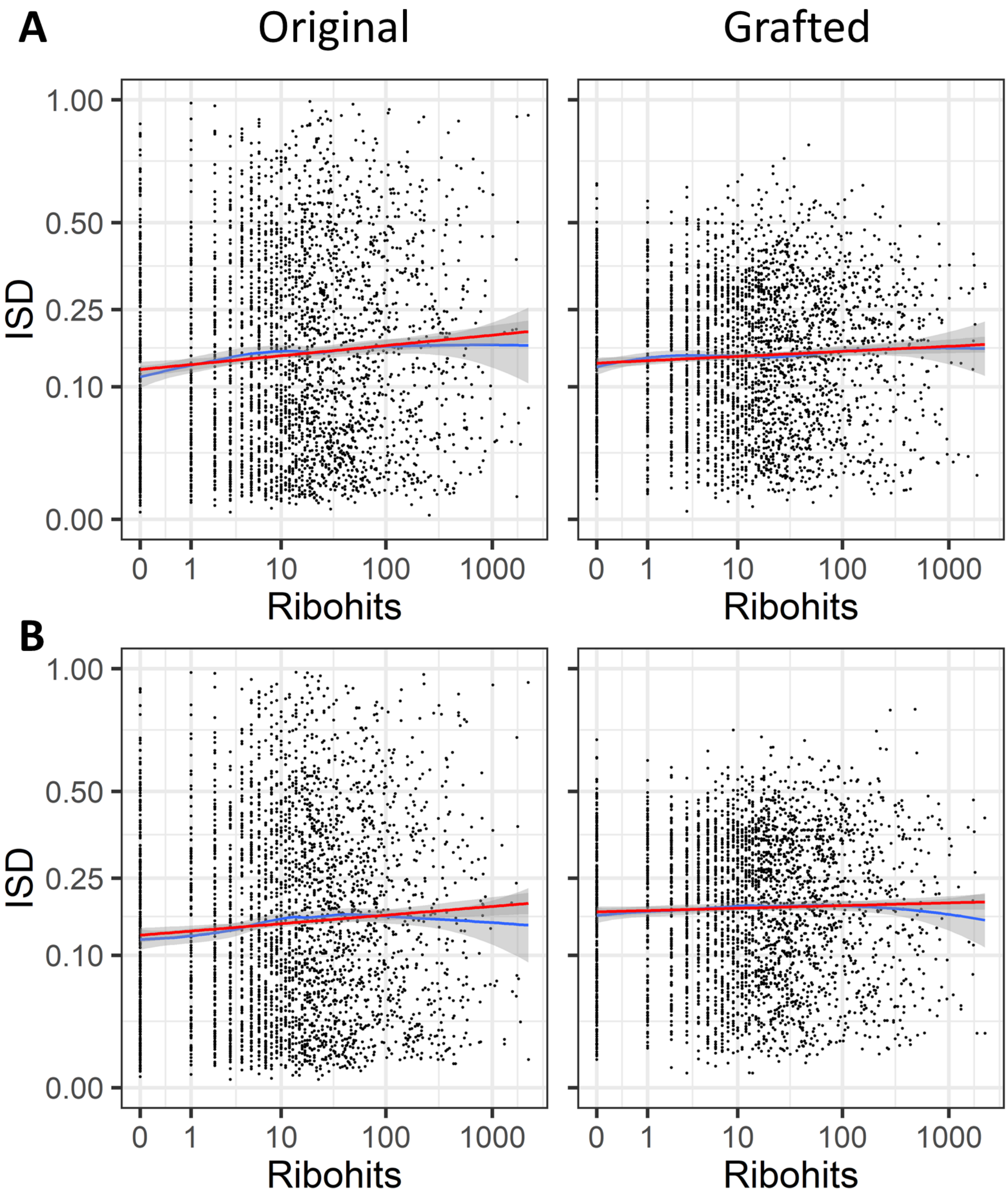


**Supplemental fig. 4.** Unweighted regressions for in-frame extension ISD on ribohits do not display the pronounced downturn past 100 ribohits seen in Fig. 5A, indicating that the downturn is due to weighting by extension length. A) In-frame and B) +2 frame regressions are the same as in fig. 5A and supplemental fig. 2A, respectively, except unweighted.

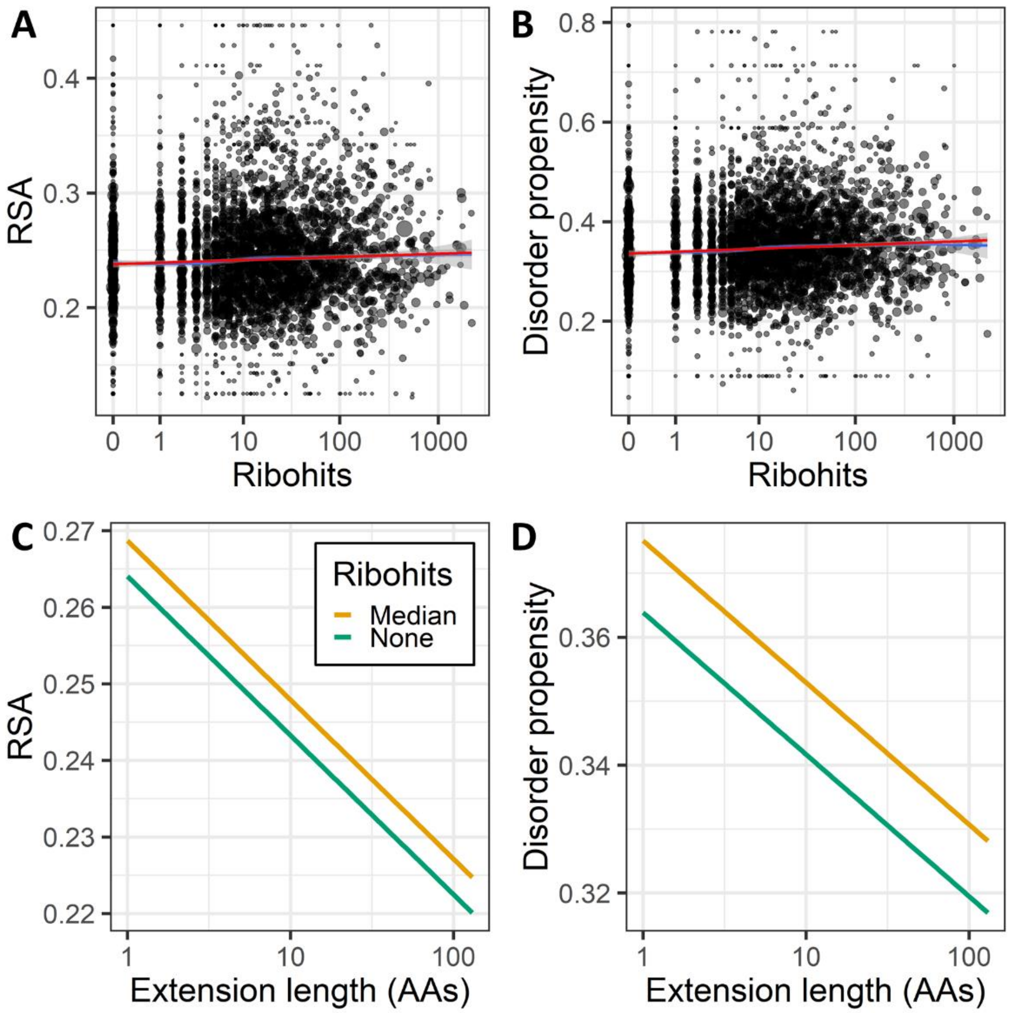


**Supplemental fig. 5.** Metrics that do not use a sliding window provide further evidence that pre-adapting selection acts on extension amino acids in +2 shifted extensions. A) Linear (red) and loess (blue) regressions of +2 frame mean RSA (left) and disorder propensity (right) on log-ribohits without controlling for length are shown. All regressions are weighted by the length of the extensions, visually represented by point area. Weighted $R^2$ = 0.0033 and 0.0061 (Willett and Singer 1988), and in the corresponding unweighted analyses $R^2$ = 0.0022 and 0.0016, for RSA and disorder propensity, respectively. B) After controlling for the effect of extension length, +2 shifted extension mean RSA and disorder propensity are still higher when ribohits are present ($P = 6 \times 10^{-5}$ and $1 \times 10^{-6}$, +2 shifted RSA and disorder propensity models, respectively, supplementary Table 3). Extension length is also predictive; +2 shifted extension mean RSA and disorder propensity are both lower for longer extensions ($P = 2 \times 10^{-24}$ and $5 \times 10^{-8}$). Weighted $R^2$ for a model with both extension length and ribohits as predictors = 0.036 and 0.015 (Willett and Singer 1988), and in the corresponding unweighted analyses $R^2$ = 0.063 and 0.026, for RSA and disorder propensity, respectively. Lines for “Median” and “None” are as described in supplemental fig. 2, except using either the RSA (left) or disorder propensity (right) model in supplemental Table 3. As in fig. 5 and supplemental fig. 2, weighting by the length of the extensions means that RSA and disorder propensity values represent expectations from sampling an amino acid from the extensions rather than from sampling an extension.

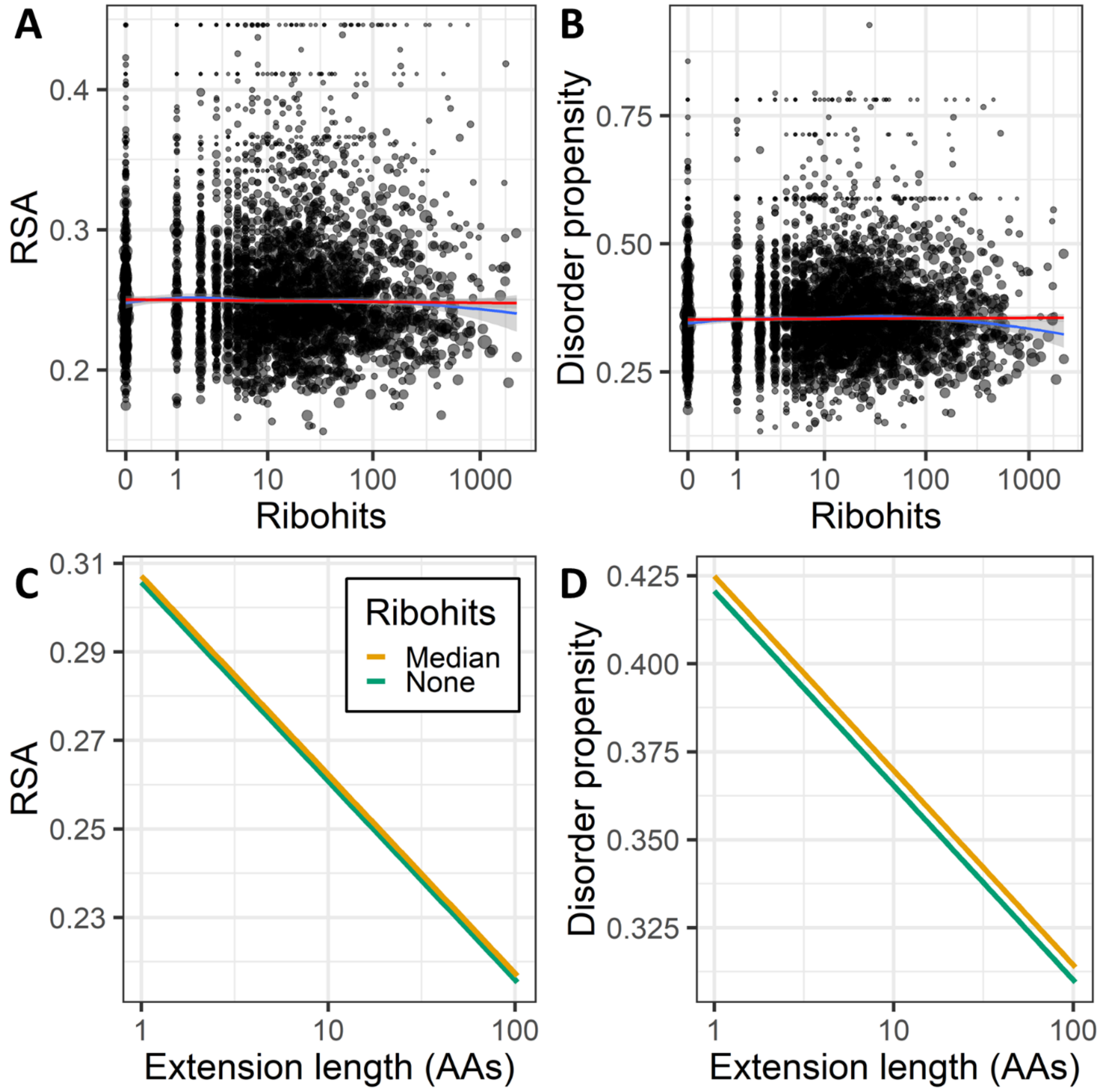


**Supplemental fig. 6.** Metrics that do not use a sliding window provide evidence that pre-adapting selection acts on extension amino acids in +1 shifted extensions via extension length but not via ribohits. A) Linear (red) and loess (blue) regressions of +2 frame mean RSA (left) and disorder propensity (right) on log-ribohits are not controlled for length. All four regressions are weighted by the length of the extensions, visually represented by point area. B) After controlling for the effect of extension length, +1 shifted extension mean RSA and disorder propensity are not significantly higher when ribohits are present ($P > 0.05$ for both +1 shifted RSA and disorder propensity models, supplementary Table 4). However, extension length is predictive; +1 shifted extension mean RSA and disorder propensity are both lower for longer extensions ($P = 4 \times 10^{-110}$ and $6 \times 10^{-40}$). Weighted $R^2$ for a model with both extension length and ribohits as predictors = 0.15 and 0.054 (Willett and Singer 1988), and in the corresponding unweighted analyses $R^2$ = 0.41 and 0.20, for RSA and disorder propensity, respectively. Lines for “Median” and “None” are as described in supplemental fig. 2, except using either the RSA (left) or disorder propensity (right) model in supplemental Table 4.

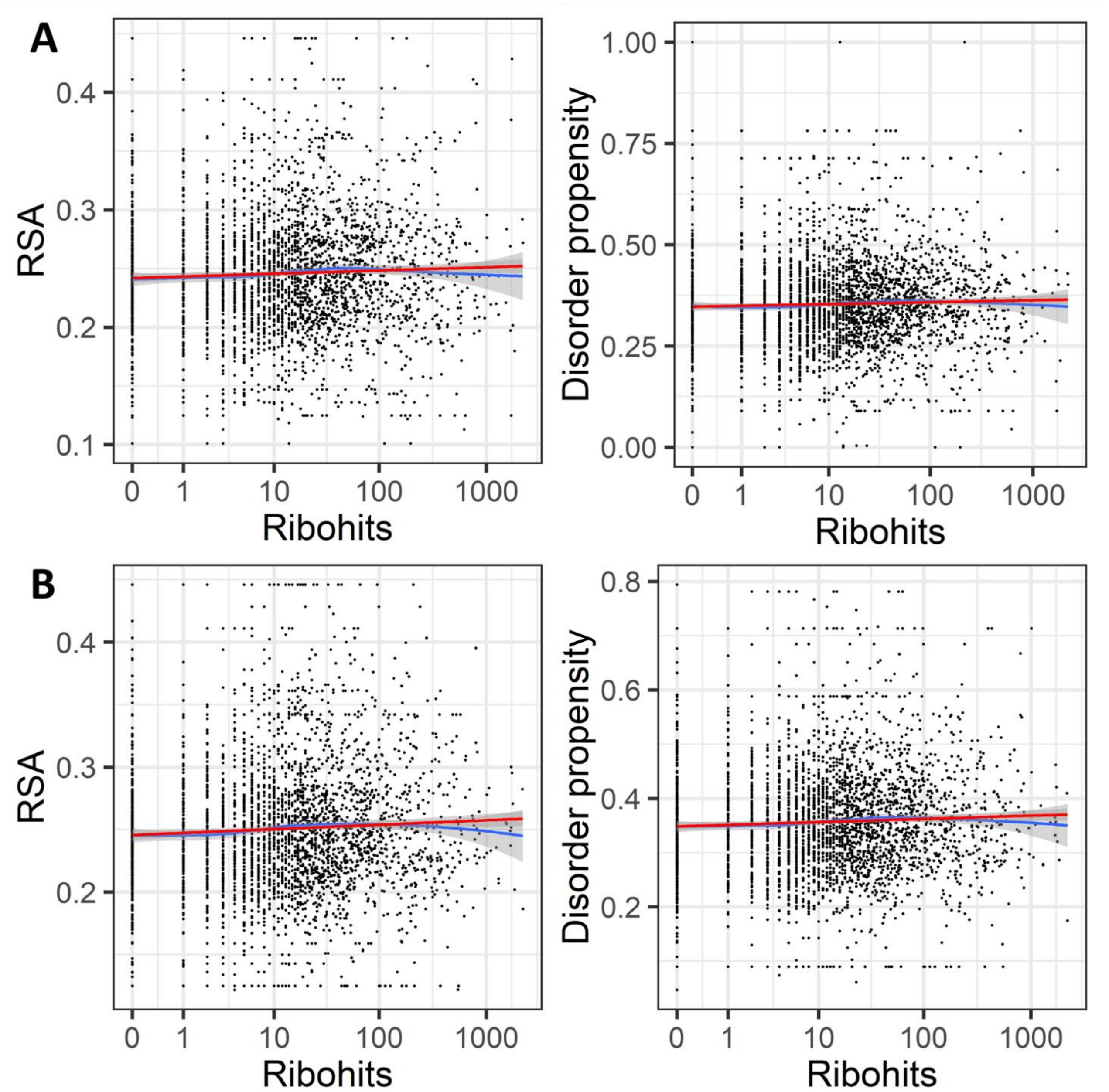


**Supplemental fig. 7.** Unweighted regressions for RSA and disorder propensity do not show a significant downturn for very large numbers of ribohits. A) In-frame and B) +2 frame regressions are the same as in fig. 6A and supplemental fig. 5A, respectively, except unweighted.

### Supplemental Tables

Methods for models in supplemental tables are the same as for the corresponding in-frame tables in the main document.

**Supplemental Table 1.** Regression model predictors of original and grafted +2 shifted ISD

| PREDICTOR FOR EXTENSION ISD | $\sqrt{Original}$ | | $\sqrt{Grafted}$ | |
|---|---|---|---|---|
| | β | P-value | β | P-value |
| $Intercept$ | 0.54 | $< 2 \times 10^{-16}$ | 0.63 | $< 2 \times 10^{-16}$ |
| $log(Ribohits + 0.5)$ | 0.019 | $2 \times 10^{-5}$ | 0.013 | $5 \times 10^{-4}$ |
| $log(Length)$ | -0.18 | $2 \times 10^{-57}$ | -0.22 | $6 \times 10^{-120}$ |

**Supplemental Table 2.** Regression model predictors of original +1 shifted ISD

| PREDICTOR FOR EXTENSION ISD | $\sqrt{Original}$ | |
|---|---|---|
| | β | P-value |
| $Intercept$ | 0.61 | $< 2 \times 10^{-16}$ |
| $log(Ribohits + 0.5)$ | 0.0065 | 0.2 |
| $log(Length)$ | -0.21 | $1 \times 10^{-73}$ |

**Supplemental Table 3.** Regression model predictors of +2 shifted mean RSA and disorder propensity.

| PREDICTOR FOR RSA/DISORDER | $RSA$ | | $Disorder\ propensity$ | |
|---|---|---|---|---|
| | β | P-value | β | P-value |
| $Intercept$ | 0.27 | $< 2 \times 10^{-16}$ | 0.37 | $< 2 \times 10^{-16}$ |
| $log(Ribohits + 0.5)$ | 0.0032 | $6 \times 10^{-5}$ | 0.0078 | $1 \times 10^{-6}$ |
| $log(Length)$ | -0.021 | $2 \times 10^{-24}$ | -0.022 | $5 \times 10^{-8}$ |

**Supplemental Table 4.** Regression model predictors of +1 shifted mean RSA and disorder propensity.

| PREDICTOR FOR RSA/DISORDER | $RSA$ | | $Disorder\ propensity$ | |
|---|---|---|---|---|
| | β | P-value | β | P-value |
| $Intercept$ | 0.31 | $< 2 \times 10^{-16}$ | 0.42 | $< 2 \times 10^{-16}$ |
| $log(Ribohits + 0.5)$ | 0.0010 | 0.2 | 0.0030 | 0.07 |
| $log(Length)$ | -0.045 | $4 \times 10^{-110}$ | -0.055 | $6 \times 10^{-40}$ |

Willett JB, Singer JD. 1988. Another cautionary note about $R^2$ – its use in weighted least-squares regression analysis. American Statistician 42:236-238.
